# Theca cell mechanosensing and regulation of follicular extracellular matrix during ovarian follicle development

**DOI:** 10.64898/2026.03.12.711479

**Authors:** Boon Heng Ng, Arikta Biswas, Kosei Tomida, Kim Whye Leong, Yuting Lou, Chin Hao Lee, Rothswell Lanting, Thong Beng Lu, Ranmadusha Merengha Hengst, Mui Hoon Nai, Isabelle Bonne, Jennifer Lauren Young, Chwee Teck Lim, Chii Jou Chan

**Author notes:** These authors contributed equally to this work (Boon Heng Ng, Arikta Biswas).

## Abstract

Mammalian folliculogenesis is essential for female hormonal regulation and successful reproduction. While the steroidogenic functions of theca cells (TCs) have been implicated in ovarian diseases and infertility, the physico-structural properties of TCs and their associated extracellular matrix (ECM), or theca matrix, remain poorly understood. Using murine ovaries, we show that a stiff basement membrane (BM) and theca matrix constitute a mechanically instructive niche that modulates TC proliferation and Yes-associated protein (YAP) signalling in secondary follicles. We identify hyaluronic acid (HA) as a key matrix component that is actively secreted by contractile TCs. The HA scaffold, in turn, regulates TC proliferation, YAP signalling and motility, and is required for overall follicle growth. We showed that stiffer substrates enhance YAP nuclear transport in TCs, while mechanical stretch, cell packing, and curvature affect TC proliferation. In addition, TCs exhibit directed migration towards regions of positive curvature. Together, this study reveals a mechanochemical feedback mechanism that establishes TC mechanics and HA as key regulators of theca matrix formation that is essential for mammalian folliculogenesis.

**SIGNIFICANCE:** The structural properties and mechanical functions of the basement membrane and theca cell-matrix encapsulating mammalian ovarian follicles are poorly understood. Our findings reveal that during early stage of follicle development, the basement membrane remains thin and stiff, while the theca cells actively secrete hyaluronic acid in a contractility-dependent manner. The hyaluronic acid scaffold, in turn, regulates theca cell YAP signalling, proliferation and motility that are required for functional growth of follicles. We further showed that the theca cells are mechanosensitive and fine-tune their proliferative capacity and Hippo pathways in response to substrate stiffness, stretch and curvature. Together, our study uncovers mechanoregulatory feedback between theca cells and associated extracellular matrix, offering new insights into environmental control of folliculogenesis in female reproduction.

## INTRODUCTION

The development of follicles within the ovary is crucial for sustaining female fertility and endocrine functions across the reproductive lifespan. During secondary follicle development, the oocyte grows in size, surrounded by increasing layers of granulosa cells (GCs) and a basement membrane (BM) that separates the follicle from the surrounding stroma. In contrast to primary follicles, an additional morphological change is the emergence of spindle-shaped TCs overlaying the BM which eventually specify into the inner theca interna and outer theca externa during antral follicle stage. The theca interna is vascularised and actively synthesises hormones and regulatory factors essential for follicle growth (1), while the externa is known to support structural integrity and ovulation (1, 2). Abnormality in TC hormonal activities has been linked to ovarian diseases such as polycystic ovarian syndrome (PCOS) and infertility (3, 4). While these studies support the functional relevance of TCs in folliculogenesis, a basic understanding of the physico-structural aspects of the TCs, its associated ECM (hereafter defined as theca matrix), and the BM remains elusive.

Emerging evidence revealed that apart from the surrounding GCs, oocyte growth is also critically influenced by the follicle’s mechano-microenvironment, such as stroma stiffness and mechanical stress(5, 6). In line with this, recent work demonstrated that in secondary follicles, the contractile TCs generate significant surface compressive stress to modulate GC YAP signalling, proliferation and overall follicle growth (7). The same study showed that the theca matrix contains fibronectin, which is actively secreted by TCs in a contractility dependent manner. This study opens several new questions: how do the TCs maintain their contractility during development? Are there other matrix components at the theca layer similarly regulated by TC mechanics? Do changes in matrix biophysical signals modulate TC mechanics and its functions?

Past studies on the theca matrix and BM remain limited and are largely based on bovine follicles. These studies revealed that the BM comprises mainly of collagen IV (8) and laminin, with their relative compositions changing during development (9, 10). While the BM provides structural and hormonal support for follicle growth (8), the physical properties of the BM remain uncharacterised. The theca matrix, where the TCs are embedded within, includes specialised ECM components such as collagens, proteoglycans, and glycosaminoglycans (11–13). Among these, hyaluronic acid (HA) has been observed to form a shell around growing follicles, with their overall content appearing to decrease during ovarian ageing (14) Despite being a major regulator of tissue hydration, viscoelasticity, and cell-matrix signalling (15, 16), the process of HA remodelling and its potential crosstalk with TCs, as well as its impact on overall follicle growth, remain poorly defined.

In this study, we hypothesise that during folliculogenesis, tissue geometric and mechanical cues can reciprocally guide theca layer functions via mechanotransduction, thereby contributing to robust follicle growth. Using a combination of molecular and biomechanical characterisation, along with *ex vivo* (follicle) and *in vitro* mechanical and chemical perturbations, we demonstrate a functional link between the mechanical status of TCs and the theca matrix, and that environmental biophysical signals such as tissue stiffness, stretch and curvature can trigger changes in TC functions. Together, our findings uncover a previously unrecognized role of TC mechanosensing in matrix secretion that contributes to the formation of theca matrix essential for functional growth of follicles.

## RESULTS

### Basal theca cells are mechanosensitive to compressive stress

We hypothesised that the TCs and GCs, located on opposite side of the curved BM, may experience distinct mechanical stress. Specifically, the elongated TCs may be subjected to tensile stress while the GCs are under compressive stress. To test this hypothesis, we immunostained murine ovarian tissue slices with YAP, a well-established mechanotransducer (17). We found that the basal TCs (TCs in contact with the BM) have higher YAP nuclear-to-cytoplasmic (N/C) ratios than the basal GCs (GCs in contact with the BM) (Fig. 1A-B). This was also observed in isolated follicles *ex vivo* (Fig. 1C-D), though their YAP N/C ratios were higher than those *in situ*. This difference may be attributed to the presence of the surrounding stroma and follicles, since tissue packing can also influence YAP activation (18). To mimic confinement *in situ*, we applied a global compressive stress (10 kPa) by incubating isolated follicles in dextran-filled medium (19). This led to a decreased YAP N/C ratios in the basal TCs (Fig. 1D), indicating that the basal TCs’ Hippo signalling activities are sensitive to surrounding tissue pressure.

**Figure 1.**
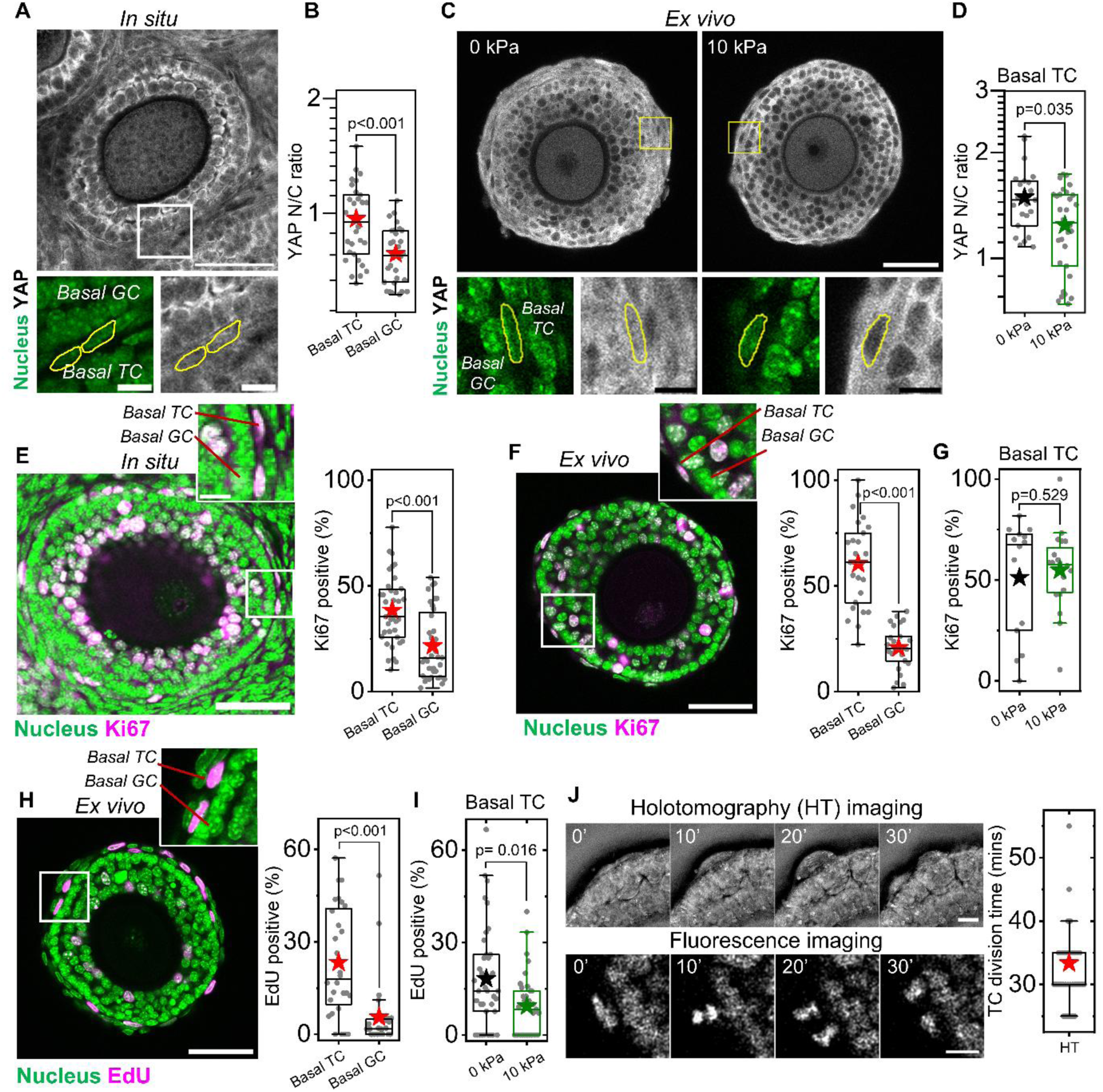
YAP nuclear transport and proliferation of basal TCs reduces on compression. (A) Representative image of an ovarian slice stained for YAP. Scale bar: 50 μm. Inset: zoomed-in region of the marked white box, stained for YAP (white) and DAPI (green). Basal GC and TC were labelled, and nuclei of basal TCs were outlined in yellow. Scale bar: 20 μm. (B) Boxplots of YAP N/C ratio in basal TCs and GCs *in situ*. *n* = 30 follicles. (C) Representative images of an isolated follicle stained for YAP in 0 kPa (left) and 10 kPa (right) conditions ex vivo. Scale bar: 50 μm. Inset: zoomed-in region of marked white boxes, stained for YAP (white) and DAPI (green) with the nuclei of basal TCs outlined in yellow. Scale bar: 20 μm. (D) Boxplots of YAP N/C ratio in basal TCs in 0 kPa (without dextran) and 10 kPa conditions *ex vivo*. *n* = 23 (0 kPa), 30 (10 kPa) follicles. (E) Left: representative image of an ovarian slice stained for DAPI (green) and Ki67 (magenta). Scale bar: 50 μm. Inset: zoomed-in region of the marked white box. Scale bar: 10 μm. Right: boxplots of proliferative basal TCs and GCs *in situ*. *n* = 39 follicles. (F) Left: representative image of an isolated follicle stained for DAPI (green) and Ki67 (magenta). Scale bar: 50 μm. Inset: zoomed-in region of the marked white box. Right: Boxplots of Ki67-positive basal TCs and GCs *ex vivo*. *n* = 46 follicles. (G) Boxplots of Ki67-positive basal TCs in follicles of 0 and 10 kPa conditions. *n* = 14 (0 kPa), 20 (10 kPa) follicles. (H) Left: representative image of an isolated follicle stained for DAPI (green) and EdU (magenta). Scale bar: 50 μm. Inset: zoomed-in region of the marked white box. Right: boxplots of EdU-positive basal TCs and GCs *ex vivo*. *n* = 31 follicles. (I) Boxplots of EdU-positive basal TCs in follicles of 0 and 10 kPa conditions. *n* = 38 (0 kPa), 33 (10 kPa) follicles. (J) Left: representative sequential still shots of basal TC division event imaged by holotomography (top) and fluorescence microscopy of H2B-labelled follicles (bottom). Numbers indicate time in minutes. Scale bar: 20 μm. Right: boxplot of TC division time measured from the holotomography images. *n* = 37 division events. Significance was determined by two-tailed Mann-Whitney U test (pairwise) in B, D-H. Boxplots show the mean (star), median (centre line), quartiles (box limits) and 1.5x interquartile range (whiskers). All data are from at least three biological replicates.

We next examined the proliferative dynamics of basal TCs. On staining for ovarian tissue slices with Ki67, a marker for active proliferation (20), we found that the basal TCs are more proliferative than the basal GCs, both *in situ* (Fig. 1E) and *ex vivo* (Fig. 1F). A similar pattern was observed when the follicles were stained with EdU (Fig. 1H), an S-phase specific proliferation marker (21). Notably, the number of EdU+ basal TCs reduced upon global compression (Fig. 1I), despite unchanged number of Ki67+ basal TCs (Fig. 1G). This likely reflects the different proliferation marker kinetics where Ki67 has a much slower turnover while EdU incorporation captures DNA synthesis within a short labelling window (20, 21). Finally, using label-free holotomography imaging (Fig. 1J, top and Movie S1), we found that the mitotic phase, spanning from cell rounding to cytokinesis. lasts approximately 35 minutes (Fig. 1J, right). A similar timescale of cell division was observed for H2B-mCherry expressing follicles (Fig. 1J, bottom and Movie S2). This timescale is relatively short compared to the mitotic duration for mammalian cells which typically lasts for an hour (22). Collectively, these data suggest that the basal TCs, in response to confinement, regulate cell proliferation, potentially downstream of mechanosensitive Hippo signalling pathway.

### Theca cells are surrounded by distinct mechanical niche in secondary follicles

We next examined changes in the TC microenvironment during secondary follicle development, focusing on the BM and theca matrix, which were rarely studied in the past. To investigate BM remodelling, we first quantified BM thickness, using high-resolution scanning electron microscopy (SEM, Fig. 2A). We found that the BM thickness remains unchanged at 45 nm in early secondary follicles (110-180 µm), followed by a steady increase and decrease again as the follicles grow past 200 µm in diameter (Fig. 2B). This reflects the highly dynamic nature of BM biogenesis, potentially to provide structural support during follicle expansion.

**Figure 2.**
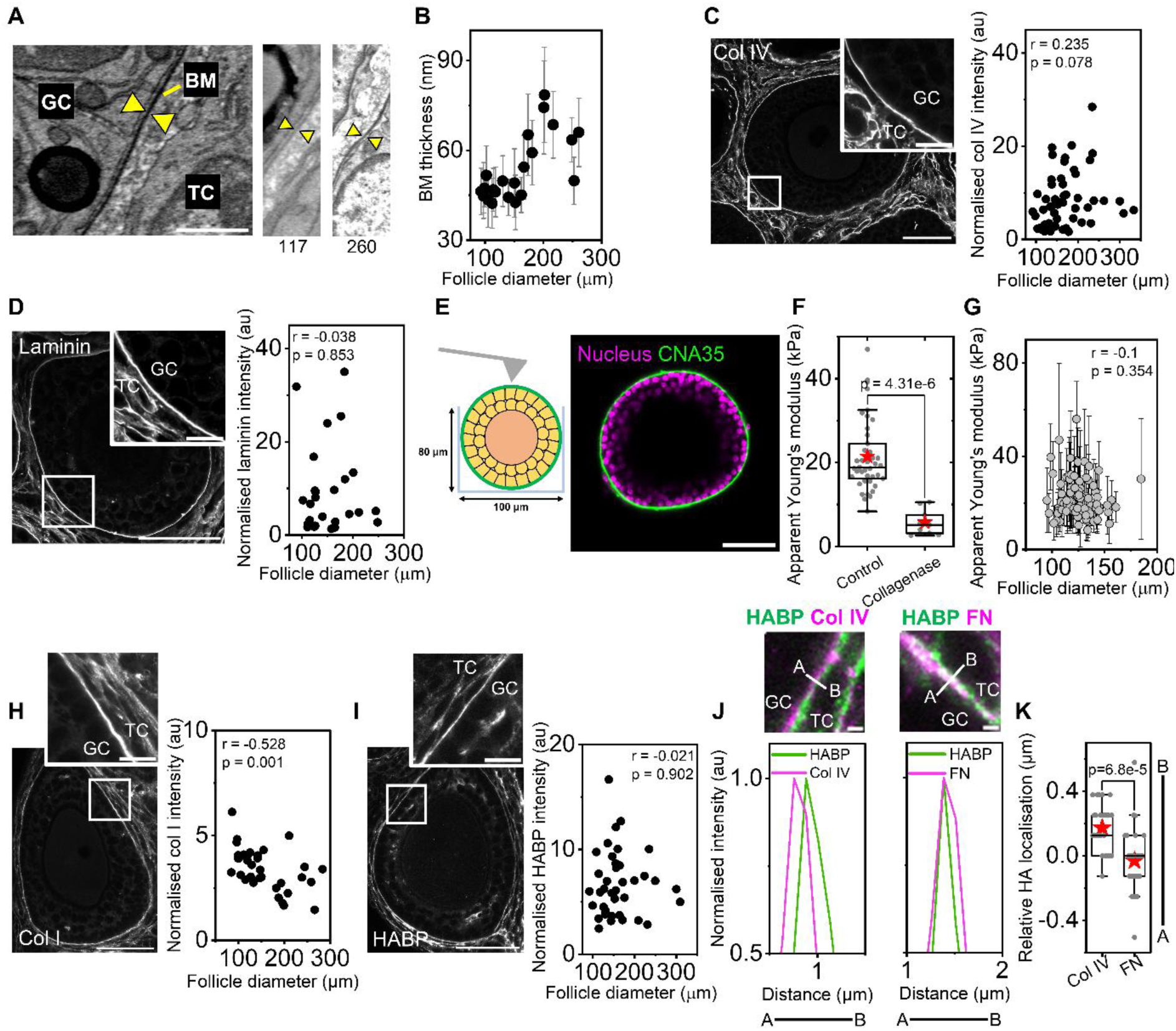
*In situ* characterisation of basement membrane (BM) and theca matrix during secondary follicle development. (A) Representative images of the BM (yellow arrowheads) via scanning electron microscopy, in early (117 μm) and late (260 μm) secondary follicles. Scale bar: 1μm. (B) Plot of BM thickness against follicle diameter. Black circles and error bars represent mean and standard deviation. *n* = 23 follicles. (C) Left: representative image of an ovarian slice stained for collagen IV. Scale bar: 50 μm. Inset: zoomed-in region marked in white. Scale bar: 10 µm. Right: plot of collagen IV intensity at BM against follicle diameter. *N* = 57 follicles. (D) Left: representative image of a tissue slice stained for laminin. Scale bar: 50 μm. Inset: zoomed-in region marked in white. Scale bar: 10 µm. Right: Plot of laminin intensity at BM against follicle diameter. *n* = 26 follicles. (E) Left: Schematic of the atomic force microscopy to measure the BM stiffness of isolated follicles. Right: representative image of an isolated follicle stained for DAPI (magenta) and CNA35 (green), after AFM measurement. Scale bar: 50 μm. (F) Plot of apparent Young’s modulus of isolated follicle BM with collagenase treatment. *n* = 41 (control), 9 (collagenase) follicles. (G) Plot of apparent Young’s modulus of the follicle BM against follicle diameter. *n* = 88 follicles. (H) Left: representative image of a tissue slice stained for collagen I. Scale bar: 50 μm. Top: zoomed-in region marked in white. Scale bar: 10 µm. Right: scatter plot of normalised collagen I intensity at the TCs against follicle diameter. *n* = 34 follicles. (I) Left: representative image of a tissue slice stained for hyaluronic acid binding protein (HABP). Scale bar: 50 μm. Inset: zoomed-in region marked in white. Scale bar: 10 µm. Right: Plot of HABP intensity at the TCs against follicle diameter. *n* = 38 follicles. (J) Top: zoomed-in images of tissue slices stained for HABP (green), collagen IV (magenta) and fibronectin (magenta). Co-localisation analysis was conducted for the line scans marked in white. Scale bar: 1 μm. Bottom: Plot of intensity profile of HABP and collagen IV or fibronectin. (K) Boxplots of HA localisation relative to collagen IV and fibronectin respectively. *n* = 9 follicles, 27 line scans. Significance was determined by two-tailed Mann-Whitney U test (pairwise) in F, K. Boxplots show the mean (star), median (centre line), quartiles (box limits) and 1.5x interquartile range (whiskers). The Pearson correlation coefficient (*r*) and significance (*p*, two-tailed test) are noted in the plots of B, C, D, G, H and I. All data are from at least three biological replicates.

To explore changes in BM composition during secondary follicle growth, we immunostained the ovarian tissue and isolated follicles with collagen IV and laminin, two principal components of BM (8). Indeed, we found collagen IV to be localised primarily at the BM and stromal compartments *in situ* (Fig. 2C, left). Focusing on the BM, we found that its expression does not change significantly during development (Fig. 2C, right; Fig. S1A). Similarly, laminin was localised specifically at the BM (Fig. 2D and Fig. S1B), with no change in expressions during follicle development *in situ* and even a slight reduction *ex vivo*. Together, these data suggest minimal change of BM components during secondary folliculogenesis.

We next investigated changes of BM elasticity during folliculogenesis, using atomic force microscopy (AFM). Secondary follicles with minimal TC coverage were isolated and positioned in polydimethylsiloxane (PDMS) microwells for AFM measurements (Fig. 2E, left). Post-indented follicles were stained with DAPI and CNA35, a collagen-binding protein, to allow for selection of ‘clean’ follicles devoid of TCs (Fig. 2E, right). Focusing on this subset of follicles, we found the apparent Young’s modulus of the BM to be around 20 kPa for secondary follicles (Fig. 2F-G). Treatment with minimal dosage of collagenase resulted in a marked reduction in apparent Young’s modulus (Fig. 2F), confirming the predominant role of collagen in BM elasticity. The minimal change in BM elasticity during early secondary follicle development (100-150 µm) is consistent with the stable expression of collagen IV and laminin in these follicles (Fig. 2C-D).

To understand if other extracellular matrix components differ across follicular compartments, we immunostained tissue slices and isolated follicles with collagen I (col I) and HA-binding protein (HABP). *In situ*, we found col I to be localised at the theca matrix and stroma. Their expression near the BM decreases with follicle size (Fig. 2H), consistent with past reports (23). In contrast, in *ex vivo* condition, col I was localised at the theca matrix and within the TCs, although its expression at the basal TCs increases with follicle size (Fig. S1C). These opposing trends suggest that the basal TCs *in situ* and *ex vivo* may regulate collagen I secretion differentially, due to distinct mechanical or biochemical environments. While HABP was also observed at the theca matrix (Fig. 2I and Fig. S1D), its intensity showed no correlation with follicle diameter *in situ* (Fig. 2I), but a slight positive correlation with follicle size *ex vivo* (Fig. S1D). To precisely determine the spatial contexts of HA, we co-stained HABP with collagen IV to indicate the BM and fibronectin (FN) to specify the theca matrix (7), respectively (Fig. 2J and Fig. S1E-F). We found that the HA scaffold resides predominantly in the theca matrix, along with fibronectin (Fig. 2K and Fig. S1G), and is spatially decoupled from the BM. Together, these data reveal that the theca matrix is enriched with HA, fibronectin and collagen I, with minimal changes of expression during secondary follicle growth.

### Contractility-mediated HA synthesis modulates TC functions

To gain further insights into HA biosynthesis at the theca matrix, we studied the existing transcriptomic datasets (24–26). By sub-clustering the theca and granulosa subpopulations of preantral follicles (named as early theca and preantral GC), along with the stromal populations in three weeks’ old mice (Fig. S2A-D), we compared HA-regulation related genes among these three clusters. Using gene set enrichment analysis (GSEA), we observed an upregulation of focal adhesions and cell-ECM receptor interactions in early TCs, which are absent in GCs (Fig. S2E). To further validate our transcriptomic profiling, we note that all theca and stroma express abundant amount of *Nr2f2*, which has been shown to be specific markers for theca cells and stroma (7, 27), but not in the GCs (Fig. 3A). Our analysis shows genes responsible for HA synthesis, particularly *Has1* and *Has2*, are enriched in the early theca and stroma, while those responsible for HA degradation, such as *Hyal2*, are expressed more in the preantral GCs (Fig. 3A). *Cemip2*, another HA regulatory gene, is also expressed at the early theca, indicative of dynamic HA remodelling in the TCs at secondary follicle stage. Finally, genes related to HA receptors, such as *Cd44* and *Hmmr*, were upregulated in theca clusters and absent in granulosa clusters. Since *Hmmr* is responsible for cell motility (28), this implies that HA-TC molecular binding may be implicated in TC motility.

**Figure 3.**
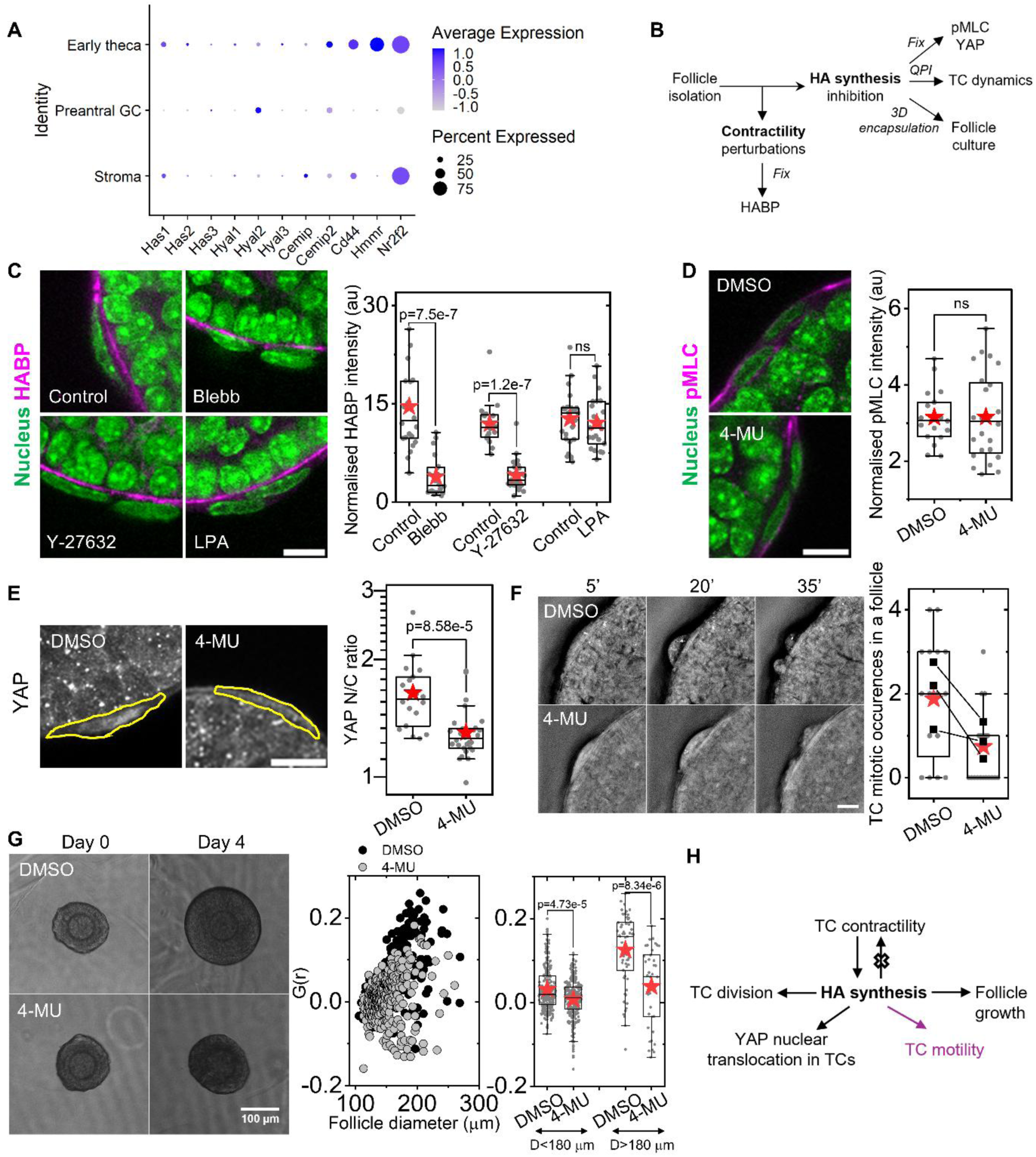
HA deposition is contractility dependent and is required for TC YAP signalling and division. (A) Transcriptomic analysis showing dot plot displaying the expression of key HA biosythesis, degradation, remodelling, and receptor-related genes in early theca, preantral granulosa, and stroma clusters in three-weeks old mice. Dot size represents the percentage of cells expressing the gene, and colour intensity indicates the average expression level. (B) Schematic depicting the interaction between TC mechanics, HA synthesis and YAP regulation. (C) Left: representative images of zoomed-in sections of isolated follicles stained for DAPI (green) and HABP (magenta) under contractility perturbations. Scale bar: 10 μm. Right: boxplots of normalised HABP intensity of basal TCs under contractility perturbations. *n* = 22 (Control), 20 (Blebb); 18 (Control), 26 (Y-27632); 24 (Control), 23 (LPA) follicles. (D) Left: representative images of zoomed-in sections of isolated follicles stained for pMLC on 4-MU treatment. Scale bar: 10 μm. Right: boxplots of normalised pMLC intensity of basal TCs on inhibiting HA synthesis. *n* = 18 (DMSO), 24 (4-MU) follicles. (E) Left: representative images of zoomed-in sections of isolated follicles stained for YAP on inhibition of HA synthesis. The outline of TC is marked in yellow. Scale bar: 10 μm. Right: boxplots of YAP N/C ratio of basal TCs on inhibition of HA synthesis. *n* = 18 (DMSO), 28 (4-MU) follicles. (F) Left: representative sequential images of basal TCs undergoing cell division on control (top) and 4-MU treated (bottom) follicle. Scale bar: 10 μm. Right: boxplots of TC mitotic occurrence in a follicle under control and 4-MU treatment. Square symbols and lines represent the average occurrences in each experiment before and after the treatment. *n* = 16 (DMSO), 19 (4-MU) follicles. (G) Left: representative images of encapsulated follicles on 4-MU treatment in day 0 and day 4 of cultures. Scale bar: 100 µm. Mid: scatter plot of growth rates per day vs. follicle size in DMSO (black) and 4-MU (grey) conditions. Right: boxplots of growth rates at pre-maturation (D<180 µm) and maturation (D>180 µm) phases in different conditions. (H) Schematic representing how HA synthesis affects theca cell properties and functions. Briefly, HA synthesis is regulated by TC contractility while disrupting HA synthesis of theca cells leads to cytoplasmic translocation of YAP, reduced division, increased motility (marked in magenta from *in vitro* experiments), and reduced follicle growth. Significance was determined by two-tailed Mann-Whitney U test (pairwise) in C-G. Boxplots show the mean (star), median (centre line), quartiles (box limits) and 1.5x interquartile range (whiskers). All data are from at least three biological replicates.

To understand how HA synthesis at the theca matrix is regulated in secondary follicles, we next investigated factors affecting HA synthesis (Fig. 3B). Since TC contractility has been shown to regulate fibronectin organisation at the theca matrix (7), we ask if TC contractility might similarly influence HA expression. Following validation of the efficacy of blebbistatin (Blebb) and Y-27632 in reducing TC contractility (Fig. S3A-B), we studied the impact of these inhibitors on HA expressions at theca matrix. Notably, reduced TC contractility led to a striking decrease of HA expressions at the theca matrix (Fig. 3C). The same trend was observed when we quantified the number of bright HA aggregates in 3D projections of follicles (Fig. S3C-D). By contrast, LPA treatment, which enhances TC contractility, does not significantly increase TCs’ pMLC expression, nor HABP intensity at the theca matrix (Fig. 3C and Fig. S3C-D). Overall, these data suggests that HA expression is mediated by TC contractility.

We next examined whether perturbing HA synthesis can in turn affect TC mechanics and functions. To confirm the efficacy of 4-MU, we first treated follicles with HA-synthesis inhibitor, 4-methylumbelliferone (4-MU), for two hours, which indeed led to a reduction in HABP intensity (Fig. S3E-F). We found that 4-MU treatment led to no change in pMLC expression (Fig. 3D), hinting minimal impact of HA on TC contractility. By contrast, 4-MU treatment led to significant reduction of YAP nuclear localisation (Fig. 3E), indicative of HA’s role in mediating TC YAP signalling. Interestingly, inhibition of HA synthesis perturbs TC proliferation and motility. Notably, follicles treated with 4-MU showed a drop in mitotic events (Fig. 3F, Movie S3), suggesting that HA regulates TC proliferative capacity. Finally, we studied the functional relevance of HA synthesis, by culturing secondary follicles with or without 4-MU (Fig. 3G). We found that 4-MU treated follicles exhibited clear attenuated growth compared to the controls, showing that an intact HA scaffold at theca matrix is essential for functional growth of follicles.

To investigate if HA secretion is driven by TCs, independent of the presence of GCs and oocytes, we isolated primary TCs from ovaries (Fig. S4A, see Methods) for cell culture studies. Isolated cells expressed high alkaline phosphatase, a marker for TCs in ovarian follicles (7), and were less circular compared to the ALP-negative GCs (Fig. S4B-C). On staining for nuclear steroid co-regulatory factor COUPTF-II (NR2F2), which is only expressed in TCs, about 67% of the cells expressed the steroidogenic marker (Fig. S4D-E), indicating reasonable TC purity. An alternative method of obtaining TCs is to harvest TCs that migrate from follicles to glass. Even though this method produces higher TC purity (95% NR2F2 positive TCs), this approach yielded substantially low number of cells for single cell studies, hence the former isolation method is chosen for subsequent studies. Since TCs are responsible for secreting androgens such as testosterone, when stimulated by human chorionic gonadotropin (hCG) (29), we next performed enzyme-linked immunosorbent assay (ELISA) to examine the testosterone levels in TC and GC cultures with or without hCG (Fig. S4F). We found that the only condition yielding a measurable level of testosterone is when TCs were cultured in the presence of hCG. Together, this suggests that isolated primary TCs retain their secretory functions post-isolation, further validating their purity.

With this approach, we confirmed that isolated primary TCs indeed secrete HA, which was depleted upon treatment with 4-MU (Fig. S5A). However, in contrast to follicle studies (Fig. 3E), YAP nuclear localisation remained unchanged with 4-MU treatment (Fig. S5B), suggesting that TC YAP signalling depends on native environmental signals. Live-cell imaging of primary TCs revealed fewer mitotic events (Fig. S5C-D), consistent with observations in follicles (Fig. 4F). Interestingly, 4-MU treatment led to increased cell motility (Fig. S5E and Movie S4), suggesting that enhanced HA synthesis impairs TC motility (Fig. 3H). Together, our results identify HA as a regulator of TC mechanobiology, linking ECM composition to cell contractility, YAP signalling, division, motility, and follicle growth (Fig. 3H).

**Figure 4.**
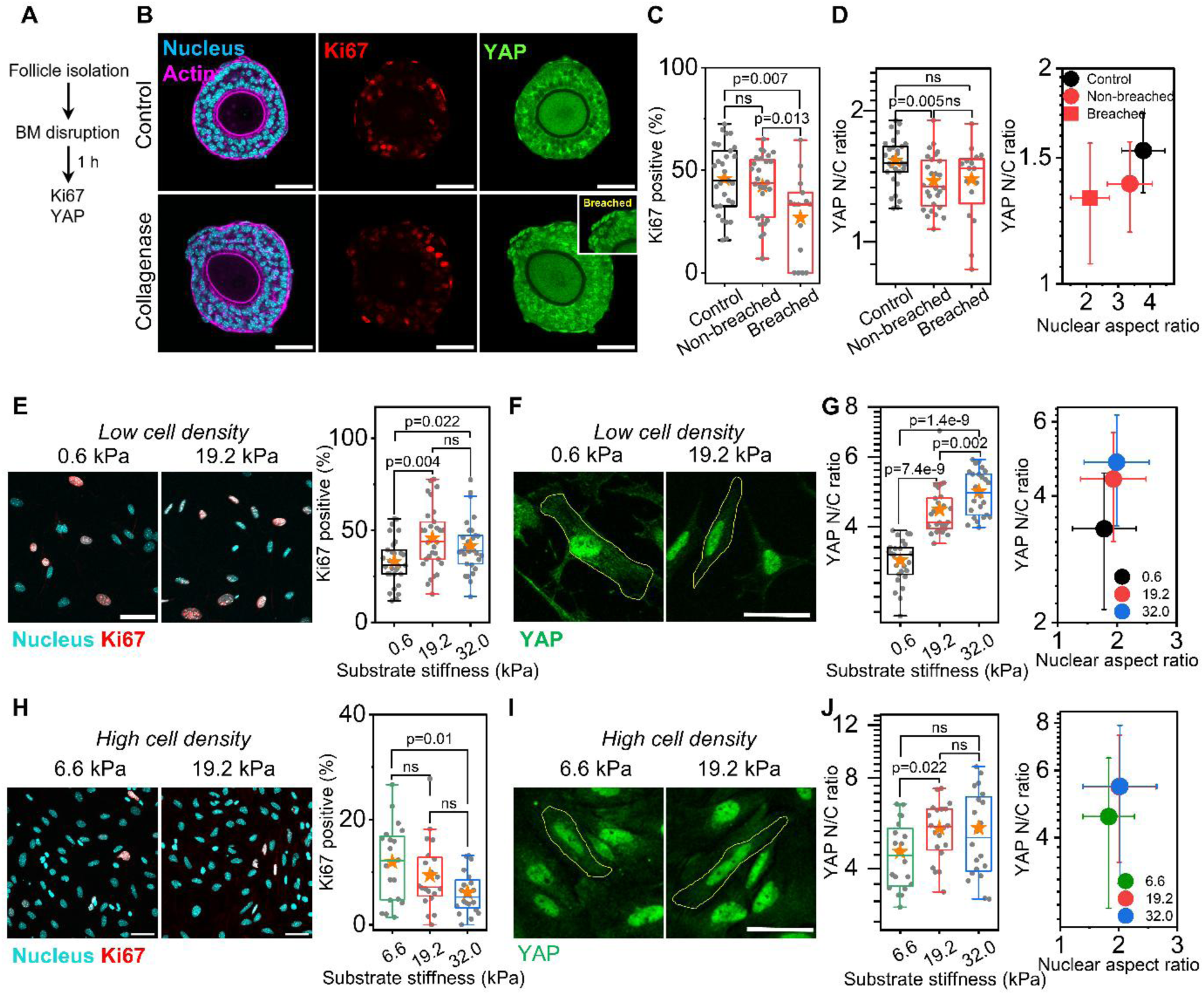
TCs are mechanosensitive to substrate stiffness and cell density. (A) Schematic of the experimental design for B-D. (B) Representative images of isolated follicles stained for DAPI (cyan), phalloidin (magenta), Ki67 (red) and YAP (green) in control and under collagenase-treatment. Scale bar: 50 μm. Inset: Yellow dotted line indicates the breached regions. of collagenase-treated follicle. (C) Boxplots of percentage of Ki67-positive TCs in control follicles and the breached and non-breached regions of collagenase-treated follicles. *n* = 29 (control), 28 (breached), 12 (non-breached) follicles. (D) Left: boxplots of YAP N/C ratio of TCs in different conditions. Right: plot of YAP N/C ratio and nuclear aspect ratio in the different conditions. Symbols and error bars represent mean and standard deviation (SD). *n* = 29 (control), 28 (breached), 12 (non-breached) follicles. (E) Left: representative images of TCs seeded at low density cultured on varying substrate stiffness and stained for DAPI (cyan) and Ki67 (red) Scale bar: 50 μm. Right: boxplots of percentage of Ki67-positive TCs cultured on varying substrate stiffness. *n* = 42 images. (F) Representative zoomed-in sections of TCs seeded at low density cultured on varying substrate stiffness and stained for YAP (green). (G) Left: boxplots of YAP N/C ratios of TCs cultured at low cell density on varying substrate stiffness. Right: plot of YAP N/C ratio and nuclear aspect ratio in the different conditions. Symbols and error bars represent mean and SD. *n* = 384 (0.6 kPa), 379 (19.2 kPa), 398 (32.0 kPa) cells. (H) Left: representative images of TCs seeded at high density cultured on varying substrate stiffness and stained for DAPI (cyan) and Ki67 (red) Scale bar: 50 μm. Right: boxplots of percentage of Ki67-positive TCs cultured on varying substrate stiffness. *n* = 20 images. (I) Representative zoomed-in sections of TCs seeded at high density cultured on varying substrate stiffness and stained for YAP (green). (J) Left: boxplots of YAP N/C ratios of TCs cultured at high cell density on varying substrate stiffness. Right: plot of YAP N/C ratio and nuclear aspect ratio in the different conditions. Symbols and error bars represent mean and SD. *n* = 200 cells. Significance was determined by two-tailed Mann-Whitney U test (pairwise) in C-E, G-H, J. Boxplots show the mean (star), median (centre line), quartiles (box limits) and 1.5x interquartile range (whiskers). All data are from at least three biological replicates.

### TCs are mechanosensitive to substrate stiffness

To further address if specific biophysical cues in the theca matrix may impact TC functions, we first tested the role of substrate stiffness, by treating isolated follicles with minimal dosage of collagenase, followed by Ki67 and YAP staining (Fig. 4A-B). Collagenase disrupted BM integrity, creating regions where the BM was compromised which we termed as breached regions. Compared to the controls, TC proliferation was reduced in the breached regions (Fig. 4C) but remained unchanged in the non-breached areas. In contrast, YAP nuclear localisation did not change significantly in the breached regions of collagenase-treated follicles (Fig. 4D, left). While we noted a reduction in the nuclear aspect ratio in TCs of collagenase-treated follicles (Fig. 4D, right), the minimal change in YAP N/C ratio indicated that nuclear elongation is not sufficient to drive YAP nuclear translocation.

To directly test TC mechanosensitivity to substrate stiffness, we cultured isolated TCs on polyacrylamide (PA) gels with stiffness of 0.6, 19.2 (similar to the BM stiffness) and 32.0 kPa, followed by Ki67 and YAP staining. The effective stiffness of the gels is validated by nano-indentation (Fig. S6A). Under sparse conditions, TC proliferation increased with substrate stiffness but became saturated above 19.2 kPa (Fig. 4E), while YAP nuclear localisation increased with stiffness up to 32 kPa (Fig. 4F-G). We observed a positive correlation between YAP nuclear translocation and nuclear aspect ratio in these conditions (Fig. 4G), suggesting that YAP nuclear entry may be regulated by force transmission to the nucleus (30). As cell proliferation and YAP signalling can be influenced by cell density (31), we tested if TCs cultured to confluency respond differently. Interestingly, we observed a significant reduction in Ki67-positive cells between 6.6 and 32.0 kPa (Fig. 4H), in contrast to the increasing trend observed in the sparse conditions. This suggested that TC proliferation may be independently regulated by cell density (Fig. 4E, H). However, YAP nuclear localisation increased consistently with substrate stiffness (Fig. 4I-J), particularly between 6.6 and 19.2 kPa, and is correlated with nuclear elongation. This indicates that substrate stiffness, rather than cell density, is a more dominant factor in influencing TC YAP mechanotransduction (Fig. 4G, J).

To determine whether substrate stiffness influences androgen production of TCs, we next cultured confluent TCs on PA gels of varying stiffnesses in the presence of hCG for three days. We found that, in the presence of hCG, TC proliferation decreased while YAP N/C ratio increased with stiffer substrates (Fig. S6B), similar to that observed without hCG. However, varying substrate stiffness did not change the testosterone secretion by TCs (Fig. S6C), suggesting that TC steroidogenic activities are not sensitive to environmental stiffness.

### TCs are sensitive to stretch and curvature

We next asked if geometric cues arising from follicle expansion can impact TC behaviour. From the growth kinetics of secondary follicles (Fig. 3G), we estimated that the follicle’s BM surface area increased by about 1% per hour, potentially causing the TCs to elongate. To determine whether TCs are mechanosensitive to stretch, we seeded primary TCs on PDMS substrates and subjected them to 4% uniaxial strain for two hours (Fig. 5A). The cells were then fixed and stained for Ki67 and YAP. We found that upon stretch, TCs displayed higher proliferation index (see Methods) compared to unstretched controls (Fig. 5B-C), though the impact on YAP localisation was minimal.

**Figure 5.**
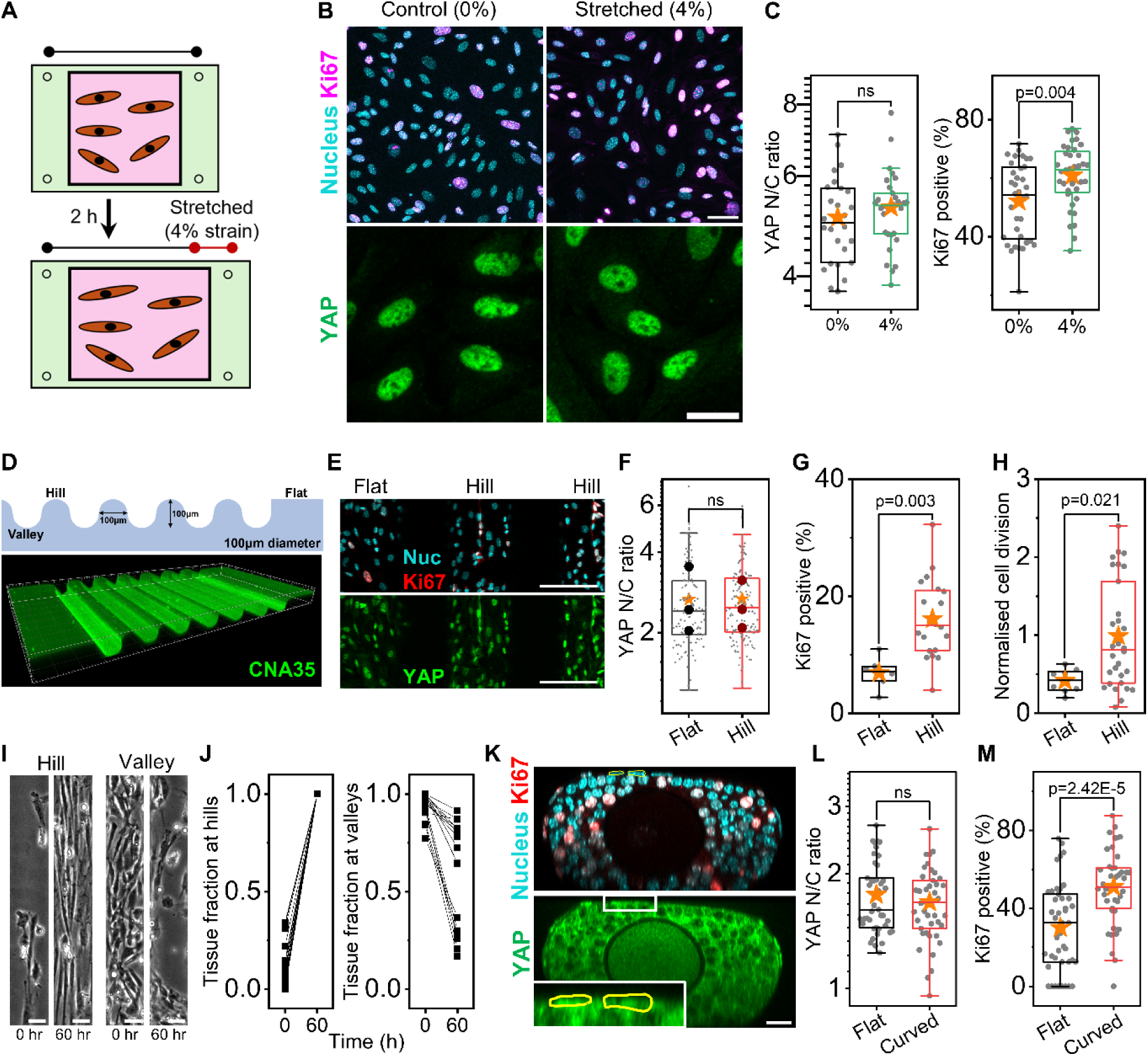
TC proliferation is sensitive to stretch and curvature. (A) Schematic showing how TCs were uniaxially stretched for 4% for 2 hours in a PDMS stretch chamber. (B) Top: representative images of isolated TCs in control versus stretched condition, stained for DAPI (cyan) and Ki67 (magenta). Scale bar: 50 μm. Bottom: representative zoomed-in images of isolated TCs in control versus stretched conditions, stained for YAP (green). Scale bar: 20 μm. Inset shows zoomed-in sections of white boxes. (C) Left: boxplots of YAP N/C ratio between control and stretched TCs. *n* = 320 (control), 400 (stretched) cells. Right: boxplots of Ki67-positive TCs in control and stretched TCs. *n* = 26 images. (D) Top: Schematic of hemicylindrical substrates with hills, valleys and flat regions. The diameter of hills and valleys is 100 μm. Bottom: 3D representative image of the substrate, coated with Col I and stained for CNA35, a collagen-binding protein. (E) Representative images of TCs at the hills and flat regions, stained for DAPI (cyan), Ki67 (red) and YAP (green). Scale bar: 100 μm. (F) Boxplots of YAP N/C ratio of TCs at various regions. *n* = 148 (hill), 121 (flat). (G) Boxplots of KI67-positive TCs at various regions. *n* = 20 (hill), 6 (flat). (H) Boxplots of normalised cell division of TCs at the hills, valleys and flats. *n* = 30 (hill), 8 (flat). (I) Representative brightfield images of TCs on the hills and valleys, 60 hours after seeding. Scale bar: 30 μm. (J) Plot of tissue fraction occupied by the TCs at the hills (left) and valleys (right), before and after 60 hours of live imaging. *n* = 18 regions. (K) Representative images of a compressed follicle stained for DAPI (cyan), Ki67 (red) and YAP (green). Scale bar: 50 μm. Inset: zoomed-in section of white box, with nuclei outlined on the top image being labelled in yellow. (L) Boxplots of YAP N/C ratios for TCs at the different regions. *n* = 43 follicles. (M) Boxplots of Ki67-positive TCs at the flat and curved regions of compressed follicles. *n* = 43 follicles. Significance was determined by two-tailed Mann-Whitney U test (pairwise) in C, F-H, L-M. Boxplots show the mean (star), median (centre line), quartiles (box limits) and 1.5x interquartile range (whiskers). All data are from at least three biological replicates.

Given the spherical geometry of the BM, we hypothesised that substrate curvature may be another physiological cue to trigger TC mechanotransduction. To test TC curvature sensing *in vitro*, we fabricated hemicylindrical arrays comprising alternating valleys, hills, and flat regions, with 100 µm radius of curvature to mimic early secondary follicle diameter (Fig. 5D). We seeded primary TCs onto these substrates, followed by immunostaining with Ki67 and YAP after 3 days of culture (Fig. 5E). Though TCs on the hills exhibited similar nuclear YAP localisation with those on the flat regions (Fig. 5F), the proliferation index is higher on the hills (Fig. 5G). Live imaging revealed that the TCs on the hills underwent more cell division than those on the flat regions (Fig. 5H), consistent with the IF quantification. We also observed that TCs demonstrated active migration towards the hills (Movie S5), eventually aligning themselves along the axis of zero curvature. In contrast, cells at the valleys moved relatively randomly (Movie S6), and by 60 hrs, the valleys were depleted of cells (Fig. 5J) while the hills were densely populated with aligned TCs.

To validate TC curvature sensing in *ex vivo* conditions, we confined isolated follicles in microwells (see Methods) to generate regions of distinct curvatures at the follicle boundary, enabling the basal TCs to assume a flat or curved geometry (Fig. 5K). Although there was no difference in the basal TCs’ YAP nuclear localisation at the flat and curved regions of the follicles (Fig. 5L), we observed a significantly higher proliferation index at the curved regions compared to the flat regions (Fig. 5M). These observations corroborate with our *in vitro* findings that while cell elongation and positive curvature are sufficient to promote TC proliferation, their impact on TC YAP signalling at this timescale remains minimal.

## DISCUSSION

Over the past decades, extensive research has established the ovary as a highly dynamic, mechano-sensitive organ (32). In contrast to the numerous studies on the ovarian stroma mechanics (14, 33, 34), the roles of TCs and follicular ECM mechanobiology remain relatively understudied. Recent work revealed that the contractile TCs help to generate intrafollicular pressure required for optimal follicle growth (7) and ovulation (2), yet the mechanical and structural properties of theca matrix, and the aspects of TC mechanosensing, remain enigmatic, partly due to challenges in physical measurements and manipulations *in vivo*. In this study, we integrated biophysical tools, chemical and mechanical perturbations with *ex vivo* and *in vitro* approaches to examine TC mechanosensing and its roles in theca matrix regulation. We demonstrated that the TCs are mechanically sensitive cells whose functions are shaped by dynamic HA synthesis and regulated by diverse mechanical cues such as stiffness, stretch, and curvature (Fig. 6).

**Figure 6.**
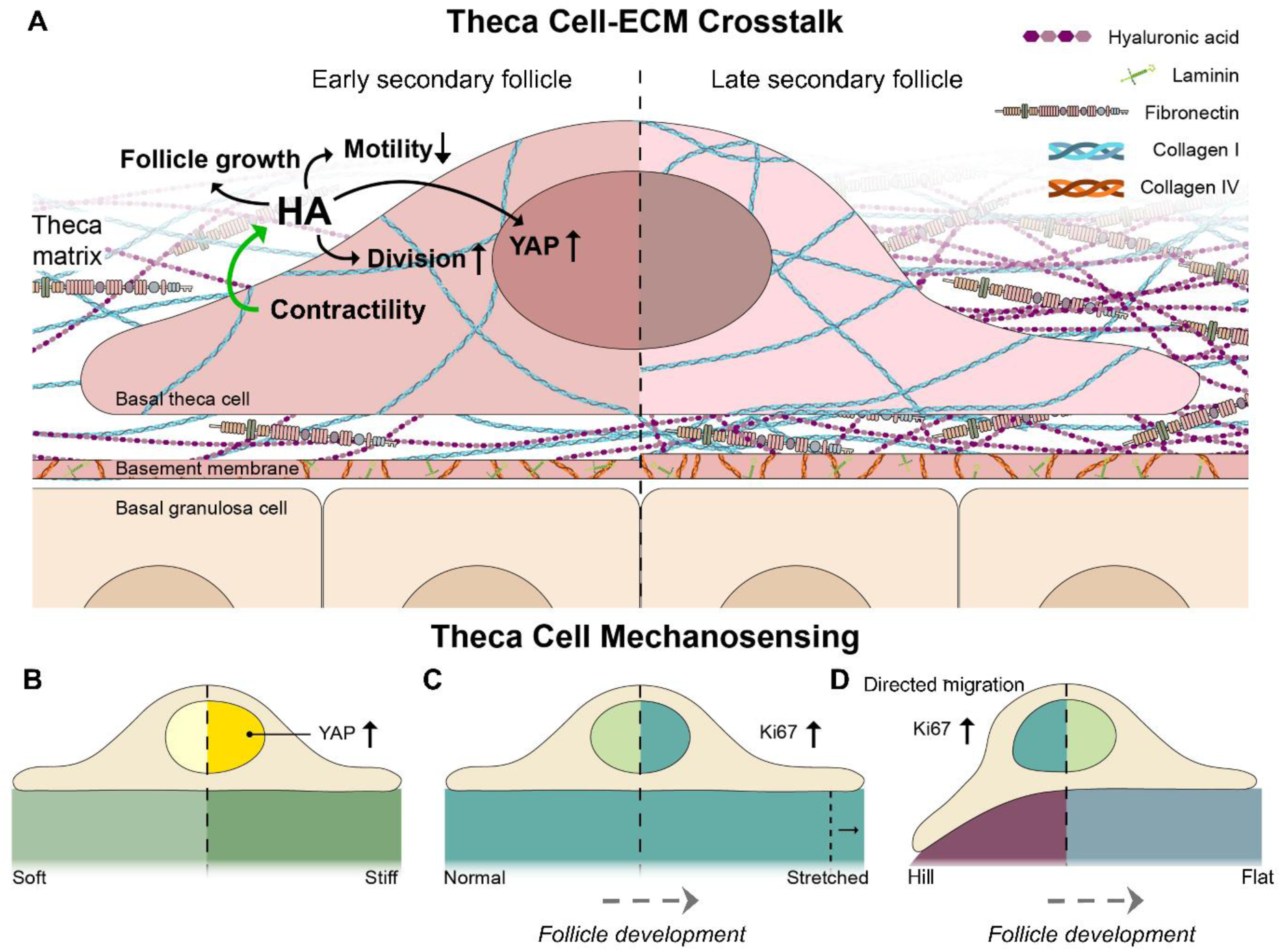
Dynamic crosstalk between mechanosensitive TCs and HA-enriched matrix regulates TC functions during secondary follicle development. (A) Schematic depicting *ex vivo* findings of increased BM thickness and various matrix depositions (Col I, fibronectin, HA) during secondary follicle growth. The maturation of theca layer involves TC-ECM crosstalk, where changes in TC contractility regulate HA expression, which in turn influences TC division, motility and YAP signalling. (B-D) Schematic showing *in vitro* and *ex vivo* evidence of TC mechanosensing. Increased substrate stiffness promotes YAP nuclear transport (B), while stretching (C) and positive curvature (D) both enhance cell proliferation. Positive curvature can additionally trigger preferential migration (curvotaxis).

Depending on the animal models and experimental approach, BM stiffness can range between 100 Pa and 10^6^ Pa (35–37). We find the BM stiffness of murine ovarian follicles lies in the range of tens of kPa, comparable to the ovarian stroma stiffness measured by AFM (14) and optical elastography methods (38). Despite its high stiffness, the mouse follicular BM (nm range) appears much thinner than the typical values (μm range) reported in other biological systems (39–41). The increase in BM thickness and constant expressions of collagen IV and laminin during secondary follicle development (100-150 µm) implies a significant volumetric growth of basal lamina, which may be actively secreted by the GCs (8). This active remodelling of BM may be functionally required to counter the increase in intrafollicular pressure during follicle development(7). Indeed, BM mechanics is critical in regulating various developmental processes, such as in *Drosophila* epithelium (42) egg chamber (43), and wing disc formation (44). Compared to the BM, the theca matrix appears much thicker (7) and dynamic during secondary follicle growth. We found collagen I to be predominantly inside TCs, and increase in expression during follicle growth *ex vivo*, similar to HA and FN synthesis (7). We note that while this trend is opposite to the decline in collagen I expressions during follicle growth *in situ*, which is consistent with observations in sheep follicles (23). These different trends may be attributed to the higher tensile stress at the surface of isolated follicles, which triggers extensive theca matrix remodelling. Together, our findings support a model where the BM and theca matrix undergo active remodelling to support tissue structural integrity during follicle growth.

Changes in BM stiffness may modulate TC signalling. Indeed, we found, both *in vitro* and *ex vivo*, that reduced substrate stiffness can lead to decreased nuclear YAP localization and proliferation of TCs. A similar effect has been observed in ovarian cancer cells seeded on stiff substrates (45). It is likely that stiffness-mediated integrin signalling (46) and focal adhesion maturation (47) may be implicated in these processes. In confluence layer, the correlation between YAP signalling and proliferation (Ki67) is broken, potentially due to TC contact inhibition, which is known to constrain cell-cycle progression (48). These findings highlight the complex interplay between substrate stiffness and cell crowding, in regulating TC behaviour (Fig. 6B).

While HA remodelling has been reported in ovarian ageing (14) and ovulation (2), its role in secondary follicle morphogenesis and interactions with TCs, is rarely studied. Here, we report a pronounced enrichment of HA at the theca matrix surrounding the follicles, consistent with a prior study (14). We further demonstrate that HA expression is directly regulated by TC contractility. Although HA secretion has been described as a mechanosensitive process (49, 50), the exact molecular pathway coupling contractile forces to HA production remains undefined. One potential candidate is ROCK signalling pathway, inhibition of which can lead to downregulation of *Has2* expression and reduced HA synthesis in fibroblasts (51), as well as altered HA-CD44 interactions in tumour cells (52). Recent studies also show that physical cues trigger specific actomyosin organisation, by balancing Rho-ROCK patterns, and directly drive metabolic changes to boost HA secretion (53). Of note, another theca matrix component, FN, was also reported to be modulated by TC contractility (7). Collectively, this evidence highlights the broad mechano-regulatory functions of TCs in theca matrix formation during ovarian follicle development.

We find that HA synthesis directly impacts YAP signalling in TCs, independent of actomyosin contractility. Such effect is less pronounced *in vitro*, suggesting that HA-YAP crosstalk in TCs requires additional matrix or GC-derived signals *in situ*. Consistent with other studies (54), we find that inhibiting HA synthesis reduces TC proliferation. However, unlike previous reports linking 4-MU treatment to reduced cell motility (54), our findings show increased TC motility upon inhibition of HA synthesis, suggesting increased cell-HA binding interactions. It is probable that reduced pericellular matrix formation (55) or weakened receptor crosstalk (56) on 4-MU treatment alters mechanical constraint, thereby facilitating migration. Another possibility arises from upregulation of matrix metalloproteinases (57) or cytoskeletal regulators from compensatory engagement of alternative ECM pathways (58), shifting towards a more migratory phenotype after transient HA loss. The elevated expression of HA receptors, *Hmmr* and *Cd44*, in early TCs, as shown in transcriptomic analysis, provides further evidence of dynamic TC-HA molecular crosstalk through mechanotransduction pathways (59–61). Given the established role of *Cd44* in mediating macrophage adhesion and migration in tissues (62, 63), a HA-rich theca matrix may influence macrophage recruitment and polarization, linking matrix mechanics to follicular development and atresia (64).

We propose that the rich scaffold of HA, fibronectin and collagen in the theca matrix, likely helps to mechanically couple the contractile mesenchymal TCs into a fluid-like tissue (65). Indeed, HA has been shown to modulate the fluidity of connective tissues and mesenchymal cells (66, 67). In recent years, HA has also emerged as a key mechanical regulator in shaping tissue morphogenesis. For example, hyaluronate pressure is shown to drive semicircular canal and cardiac valve formation (68). In the context of follicle development, the hydrated and viscous nature of HA (16) likely confers a mechanically resilient follicle shell to buffer the follicles against excessive deformation, or to serve follicle mechanosensing. HA signalling is usually dependent on its molecular weight, with high molecular weight HA being generally associated with anti-inflammatory response and tissue homeostasis (69), while fragmented, low molecular weight HA promotes inflammatory and pro-migratory signalling (70, 71). Though we did not resolve the molecular weights of HA in the theca matrix, we propose that the HA may function as a signalling hub to store various growth factors and vascular-derived cues essential for follicle growth.

To the best of our knowledge, our report is the first to demonstrate that HA is required for functional growth of follicles in mice. Our results echo those observed in rats (72), and while *Has1* gene is expressed in theca of porcine antral follicles (73) and rat preantral follicles (72), we find that *Has1* and *Has2* are primarily responsible for HA synthesis in TCs of murine preantral follicles. While the exact mechanism underlying reduced follicle growth upon 4-MU treatment remains unaddressed, we hypothesise that this may be due to attenuated compressive stress (7) or signalling roles of TCs. Future studies on transcriptomic profiling of these follicles under 4-MU perturbations, along with live imaging of intrafollicular dynamics, will further elucidate the role of HA synthesis on GC proliferation and death.

We showed that tissue geometric cues, such as substrate elongation and convex curvature, can promote TC proliferation (Fig. 6C-D). These extrinsic signals, along with matrix stiffness, are ever present and undergo constant changes during follicle development. This may potentially contribute to the morphogenesis of multi-layered theca layer. The lack of curvature-dependent YAP responses, despite its pronounced effect on proliferation, could arise from the different timescales of signalling. While YAP activity can change rapidly, Ki67 labels cell cycle progression over longer periods, hence direct comparisons of their expressions under a single experimental timeframe may not be appropriate. The decoupling between the two cellular responses also implies that TC proliferation can be controlled by geometric cues independently of the canonical YAP-mediated transcription, pointing to additional mechanotransduction pathways that warrant further exploration. Our findings contrast with that of past studies showing that positive curvatures suppress cell proliferation (74), suggesting that the observed curvature-induced proliferation may be unique to TC identity, and likely depend on other factors such as cell density and ECM coating. Similarly, our observation of TC curvotaxis towards the hill, followed by cell alignment along the zero curvature, differs from that of T-lymphocytes, mesenchymal stem cells, and fibroblasts, which were reported to migrate faster on concave surfaces (75–77). Exploring the molecular mechanisms underlying TC curvotaxis and stretch sensing constitute exciting aspects for future studies.

In summary, we report a previously unrecognized role of TC mechanosensing in the context of ovarian follicle development. The study reinforces the notion that cross-scale feedback interactions between tissue geometry, mechanics and cellular behaviour underlies robust tissue morphogenesis (78, 79). Beyond advancing fundamental ovarian biology, our findings also provide a new framework in understanding ovarian ageing and diseases. For example, age-related dysregulation of HA synthesis and turnover (80), together with increased matrix fragmentation (81) and stromal stiffening (14), are known to alter tissue viscoelasticity, which may impair TC functions and follicle growth. Finally, our work have implications for future clinical treatments of reproductive disorders, where physiologically relevant ECM components and biophysical cues, as revealed in the present study, maybe incorporated to improve fertility outcome.

## ACKNOWLEDGEMENTS

We thank Zihao Wu and Nicole Loh Kit Ying for the initial optimisation of primary cell isolation. We thank Cheng-Kuang Huang for providing the hemicylindrical moulds, and Dang Hairuo (Ellenberg group, EMBL Heidelberg) for providing ovarian tissues from 48 weeks of R26-H2B-mCherry mice (Identifier: CDB0239K). We are grateful to Lucie Kim for the illustration, the MBI Core facilities for their continuous support, the Singapore Microscopy and Bioimaging Analysis (SiMBA, MBI) core for microscopy, and the Microfabrication core for help with the microwells. The Chan lab is supported by the Ministry of Education under the Research Centres of Excellence programme through the Mechanobiology Institute and the Department of Biological Sciences at the National University of Singapore, the Ministry of Education Tier2 grant (T2EP30222-0026) National Research Foundation Mid-size Grant (NRF-MSG-2023-0001) and the Bia-Echo Asia Centre for Reproductive Longevity and Equality (ACRLE) at the National University of Singapore. C.J.C. acknowledges the support of the Singaporean Teaching and Academic Research Talent Inauguration Grant (START). We thank Neha Paddillaya and Huan Ting Ong for providing valuable feedback to our manuscript.

## AUTHOR CONTRIBUTIONS

Project Conceptualization and Design: B.H.N., A. B., C.J.C.; Experiments: B.H.N., A.B., K.T., K.W.L., C.H.L., R.L., T.B.L., R.M.H.; Data Analysis, Quantification and Statistical Analysis: B.H.N., A.B., K.W.L., Y.L., C.H.L., R.L., R.M.H., C.J.C.; Writing: B.H.N., A.B., C.J.C.; Data Interpretation: B.H.N., A.B., K.W.L., Y.L., T.B.L., I.B., C.J.C.; Supervision: C.J.C.

## METHODS

### Animal handling and maintenance

C57BL/6NTac female mice, aged P25-P28, were grouped housed in individually ventilated cages with access to water and food under a 12-hour light/12-hour dark cycle. Mouse rooms were maintained at 18-25℃ and 30-70% relative humidity. The mice were euthanised by carbon dioxide asphyxiation followed by cervical dislocation. Ovaries were then dissected from the mice and transferred to an isolation buffer consisting of Leibovitz’s L15 medium (21083027, Thermo Fisher, USA) supplemented with 3 mg/ml bovine serum albumin (BSA, A9647, Sigma, USA). All animal work was approved by the Institutional Animal Care and Use Committee (IACUC) at the National University of Singapore (NUS). R26-H2B-mCherry female mice, aged 48 weeks, were bred and maintained by Ellenburg’s Lab in EMBL (European Molecular Biology Laboratory).

### Tissue sectioning

Ovaries were dissected and cleaned in isolation buffer before fixing in 4% paraformaldehyde (PFA, sc-281692, Santa Cruz Biotechnology) at room temperature for an hour. The fixed ovaries were washed in phosphate-buffered saline (PBS) thrice before being embedded in 4% low-melting point agarose (16520050, Thermo Fisher). The embedded tissue was sliced into 100 µm tissue sections using a vibratome (VT1200S, Leica Microsystems, Germany) at a speed of 0.05 mm/s and an amplitude of 1 mm in phosphate buffer saline (PBS, Gibco).

### Follicle isolation

Follicles were mechanically isolated from dissected ovaries under the stereomicroscope attached to a thermal plate, in isolation buffer, using tweezers at 37℃. Growth medium, consisting of MEMα GlutaMAX (Thermo Fisher) supplemented with 10% fetal bovine serum (FBS, Thermo Fisher), 1% Penicillin-Streptomycin (Pen-Strep, Thermo Fisher), 1x Insulin-Transferrin-Selenium (Thermo Fisher), and 50 mIU/ml follicle-stimulating hormone (Sigma), was prepared. Individual follicles were transferred to a 96-well non-treated plate and cultured in the growth medium at 37℃, 5% CO_2_, and 95% humidity overnight.

### Pharmacological treatments

Isolated follicles and TCs were treated with 170 μM 4-methylumbelliferone (4-MU, Sigma) to perturb HA synthesis for 2 hours. To investigate how HA synthesis inhibition impacts TC division and motility, the same isolated follicles and TCs were initially cultured as the control condition before treating with 170 μM 4-MU for at least 3 hours. Isolated follicles were treated with either 20 μM blebbistatin (Selleck) or Y-27632 (Merck) for 2 hours to inhibit contractility and 20 μM LPA (Oleoyl-L-a-lysophosphatidic acid, Sigma) for 2 hours to enhance contractility. Follicles were treated with 1 mg/mL collagenase IV (Gibco^TM^) for an hour to perturb basement membrane stiffness for the validation of the AFM measurement; and with 0.1 mg/mL collagenase IV for 2 hours to assess the transient effect of BM integrity perturbation on TC functions.

### Quantitative phase microscopy

The microwell mould was 3D-printed with resin, with a diameter of 2 mm and a depth of 2 mm. PDMS microwells were then fabricated using the moulds to house isolated follicles (82). The microwells were mounted onto a plastic dish and sterilised with UV for at least 15 minutes. 1% alginate (Sigma) was prepared in PBS and further mixed 1:1 with growth medium. The 0.5% alginate solution was then added to the microwells. Crosslinking medium, comprising 50 mM calcium chloride (Sigma) and 140 mM sodium chloride (1^st^ BASE), was added after the follicles were transferred to the microwells. After 2-3 minutes, the leftover crosslinking medium was carefully removed before supplementing with more growth medium. The dish was then mounted and equilibrated on the vessel holder of the Tomocube (HT-X1), controlled at 37℃ with 5% CO_2_. After determining minimal drift from the encapsulated follicles, holotomograms of the follicles were obtained using a low-coherence HT system (HT-X1, Tomocube Inc., Korea). HT-X1 employs an incoherent 450-nm LED light for illumination, and the follicles were imaged at a z step of 0.78 μm over a range of about 142 μm, with a time step of 5 minutes. For the experiment, the follicles were first imaged for 2-3 hours as the control condition. 170 μM of 4-MU was then added to the growth medium, and the follicles were imaged for another 2-3 hours. The holotomograms were processed using TomoAnalysis.

### Live imaging of H2B dynamics

Follicles isolated from the ovaries of the R26-H2B-mCherry female mice were mounted on the Zeiss LSM880. The follicles were imaged with the non-descanned two-photon mode at the 1080 nm wavelength using the LD C-Apochromat 40x/1.1 W Corr M27, with a 2 μm z step and a 10-minute interval, for a total duration of 16 hours.

### 3D follicle culture

1% alginate (Sigma, 180947) was prepared in PBS (Gibco, 18912014) and mixed with growth medium in a 1:1 ratio. A crosslinker solution was prepared, consisting of 50 mM calcium chloride (Sigma, C1016) and 140 mM sodium chloride (1^st^ BASE, BIO-1111). Follicles were first mouth-pipetted to the 0.5% alginate solution, and hydrogels were formed by pipetting each follicle-alginate mix into the crosslinking solution for 2 mins. Once encapsulated, each gel was placed in 100 µL growth medium inside individual wells of Ultra-Low Attachment 96-well plate (Corning, CLS3474) at 37 °C, 5% CO_2_, and 95% humidity for up to 4 days. For long term cultures, half the volume of the growth medium was changed in each well every 2 days.

### Primary theca cell isolation

Primary theca cells were isolated based on the protocols adapted from Tingen et al (83). Freshly isolated ovaries were washed thrice in cold isolation buffer and poked with a needle to burst the follicles and release the GCs. The ovaries were punctured until intact follicular architecture was not visible. The remaining tissue was washed twice with isolation buffer and transferred to the digestion buffer, comprising 0.05 mg/mL activated DNase I (D4263, Sigma) and 10 mg/mL collagenase IV (17104-019, Thermo Fisher) in M199 (11150067, Gibco) medium. DNase was first activated by mixing DNase I with HBSS (14025-092, Gibco) in a 1:1 ratio. The tissue was incubated at 37℃ for an hour with gentle mixing every 15 minutes using a pipette. Once completely dissolved, the solution was centrifuged at 94×g for 5 minutes. The pellet was washed and resuspended in McCoy’s medium (16600082, Gibco) supplemented with 5% FBS and 1% Pen-Strep to obtain the theca/stroma cells. The remaining isolation medium was centrifuged at 94×g for 5 minutes, washed and resuspended in the supplemented McCoy’s medium to obtain the granulosa cells. Both cells were then seeded onto 6-well cell culture dishes (ThermoFisher Scientific, Nunclon^TM^ Delta Surface and incubated at 37℃, 5% CO_2_ and 95% humidity overnight before further experiments.

Follicles were allowed to settle on glass-bottomed dishes and incubated for 16 hours at 37 °C, 5% CO_2_, 95% humidity to ensure theca cells migrated from the follicles onto the glass and follicles did not rupture during incubation. The tissues were then carefully removed using a mouth-pipette; and the cells remaining on the glass were fixed and immunostained to compare the purity of the mechanically isolated theca cells versus TCs extracted from this method.

### Alkaline phosphatase staining

Isolated TCs and GCs were cultured on glass coverslips for 2 days before staining with the alkaline phosphatase staining kit (Abcam, ab242286). The cells were washed with PBST before fixation with the Fixing Solution for 2 minutes. The samples were then washed with PBST once before incubating in the AP Staining Solution at room temperature for 15-30 minutes in the dark. Samples were washed with PBS once before mounting onto glass slides with mounting medium at room temperature overnight. Samples were mounted and imaged with the IX83 Olympus inverted microscope.

### Scanning electron microscopy

Ovaries were fixed with 3% PFA and 2% glutaraldehyde (GA) overnight at 4℃. They were washed thrice with PBS for 5 minutes each. The ovaries were either processed as a whole organ or 150 μm tissue slices using the vibratome. The samples were incubated in 1% osmium tetraoxide (OsO4) with 1.5% potassium ferrocyanide in PBS for 1 hour on ice and then washed thrice with distilled water for 5 minutes each. The samples were then placed into 1% thiocarbohydrazide in distilled water for 20 minutes at room temperature and washed thrice with distilled water for 5 minutes each. The samples were then placed into 1% OsO4 in distilled water for 30 minutes at room temperature and washed thrice with distilled water for 5 minutes each. They were next incubated with 1% uranyl acetate in distilled water overnight at 4℃ and washed thrice with distilled water for 5 minutes each. 0.02 M lead nitrate and 0.03 M aspartic acid were mixed well together, and the pH was adjusted to 5.5. The samples were kept in lead aspartate solution for 30 minutes at 60℃ in the oven and again washed thrice with distilled water for 5 minutes each. Tissues were dehydrated with ethanol, increasing gradually from 25%, 50%, 75%, 95% and 100%, with 10 minutes in each solution on ice before washing with acetone twice for 10 minutes each on ice. For resin infiltration, the samples were placed in a 1:1 acetone-araldite resin mixture for 30 minutes and then 1:6 mixture overnight. They were then transferred to pure araldite for 1 hour in a 45℃ oven. This was done thrice before they were transferred into an embedding mould with pure araldite and cured for 24 hours at 60℃.

The embedded samples were then sectioned using a Diatome diamond knife with the Leica UC6 ultramicrotome, and 100 nm ultrathin sections were collected onto silicon wafers. SEM Imaging was done with Thermofisher FEI Quanta 650 FEG-SEM, where large area montage scans were acquired with MAPS 2.1 software using the backscatter mode (vCD detector) at 5 kV, 5 mm working distance (WD).

### Atomic force microscopy

Atomic force microscopy (AFM) was performed to quantify BM mechanical properties. The NanoWizard 4 BioScience (JPK Instruments AG, Germany), mounted on an inverted microscope (Olympus IX81) with a 10× (NA 0.30) objective, was used. As the follicles were spherical and challenging for AFM measurement on flat surfaces, multiple microwell patterns were customised with specific widths and heights that can house the follicles during AFM. The widths of the wells range from 100-150 μm and 170-220 μm. The width of the microwells was designed to ensure that follicles of different sizes could fit in the respective microwells. The height and spacing between the wells remained constant at 80/150 μm and 50 μm, respectively. The height was designed to allow the top portion of the follicles to be exposed to the AFM probe, which is critical to prevent the probe from touching the walls of the microwells and breaking.

PDMS (Sylgard 184 silicone Elastomer Base) and crosslinker (Sylgard 184 Elastomer curing agent) were mixed in a ratio of 10:1 and degassed before transferring to the wafer. The PDMS mixture was degassed again and cured at 80℃ for 2 hours. The PDMS mould was removed and trimmed to the working size. To fabricate the microwells, the PDMS mixture was transferred onto the glass bottom dish (WPI FD35), and the trimmed PDMS mould was placed inverted on top. The assembly was cured at 80℃ for 2 hours. The PDMS mould was then removed, and the PDMS microwells were used for AFM measurements. The microwells were filled with growth medium, degassed and incubated at 37℃ with 5% CO_2_ for at least 30 minutes. Freshly isolated follicles were transferred onto the microwells and left to stabilise before the AFM measurements.

To measure BM elasticity, Bruker MLCT-D cantilevers with a pyramidal tip and a nominal spring constant of 0.03 N/m were used. The sensitivity was calibrated using the contact-based approach for each experiment. BM elasticity was measured with a constant speed of 5 μm/s, with a loading force of 3 nN in a 5 μm x 5 μm area. The follicles were imaged at 10×, and the brightfield images were used to determine the follicle diameter.

### Nanoindentation

PA gels without collagen I coating were used to verify the substrate stiffness. The Young’s modulus of the PA gels was measured with a Chiaro Nanoindenter (Optics11Life, Amsterdam, The Netherlands). Hydrogels were polymerised on circular glass coverslips and swollen overnight in Milli-Q water at 4℃. Before indentation, the coverslips were affixed to a glass petri dish using super glue, and the hydrogels were immersed in Milli-Q containing 0.5% bovine serum albumin (BSA) to minimise probe-sample adhesion. The probe was selected to have a large spherical tip radius of 51 μm and a suitable cantilever spring constant (0.47 N/m). Each hydrogel surface was indented with an 8-by-8 matrix scan, spaced with a step size of 1000 μm. Indentations were performed at an approach velocity of 1.5 μm/s to a maximum depth of 3 μm. The probe was held at this depth for 1 second before being retracted over 2 seconds.

### Substrate stiffness assay

Various stiffnesses of PA hydrogels were made following specific combinations of acrylamide and bis-acrylamide (84). Glass coverslips were cleaned using the UV ProCleaner before being treated with 0.3% acetic acid and 0.5% 3-(trimethoxysilyl)propyl methacrylate (TMSPMA) in ethanol. The coverslips were washed twice with 100% ethanol and then dried. Ammonium persulfate (APS) and tetramethylethylenediamine (TEMED) were added to the PA mixture before adding onto a dichlorodimethylsilane (DCDMS)-treated glass slide and then placing the treated coverslip onto the drop of PA mixture. The coverslip, with the polymerised PA hydrogel, was removed and washed twice with water and then in 20 mM acetic acid for 30 minutes. Sulfosuccinimidyl 6-(4-azido-2-nitrophenylamino) hexanoate (Sulfo-SANPAH, 803332, Sigma), in 20 mM acetic acid, was then added and activated using the UV-LED system (UV-KUB 9) with 6% power for 4 minutes. The PA hydrogel was washed thrice with 20 mM acetic acid and twice with PBS. 0.0875 mg/mL of collagen I (bovine tail, A1064401, Gibco) in PBS was added to the hydrogels and incubated for at least 3 hours at room temperature or overnight at 4℃. The hydrogels were washed thrice with PBS before use or storage at 4℃. The PA hydrogels were first sterilised with UV light before being used to culture TCs, which were then fixed with 4% PFA.

### Stretch assay

The PDMS chamber (Strex, STB-CH-10) was sterilised using UV light a day before cell seeding. ECM protein (fibronectin, bovine) was added and incubated overnight at 4℃ before washing with PBS and culture medium. Isolated TCs were seeded onto the PDMS chambers and cultured for two days. The TCs were subjected to a 4% strain for 2 hours using the manual cell stretching system (Strex, STB-100) before fixation.

### Curvature assays

The original fabrication of the mould for the hemi-cylindrical wave substrate was described in Huang et. al (85). With the original NOA73 (Norland optical adhesive 73) master mould, multiple clones of PDMS and NOA73 moulds were fabricated. The PDMS base and crosslinker (SYLGARD^TM^ 184 Silicone Elastomer Kit, DOW) were prepared in a ratio of 10:1 and lightly poured over the NOA73 master mould and incubated at 80℃ for 2 hours. The PDMS moulds can then be peeled away from the NOA73 master mould. To fabricate more NOA73 moulds, the PDMS mould was first placed on a clean glass slide as a support. NOA73 was added to the side of the hemi-cylindrical wave pattern, and another clean glass slide was then placed on top of the NOA73 and PDMS mould. The NOA73 was crosslinked using the UV-LED system (UV-KUB 9) with 15% power for 2 minutes. The PDMS mould was peeled away from the new NOA73 mould, and its structural integrity was checked using a stereoscope. Using the NOA73 mould with a half period of around 100 μm, PA hydrogels with the hemi-cylindrical wave pattern were cast with TMSPMA-treated coverslips. The silicon wafer for the dome structure with an estimated diameter of 80 μm was designed and fabricated by the microfabrication core facility in the Mechanobiology Institute, NUS.

PA substrates were then coated with collagen I as described previously. The PA substrates were sterilised with UV light before seeding and culturing the isolated TCs. The coverslips were mounted on coverslip cell chambers (SC15022, Aireka Cells) and the TCs were imaged live with BioStation IM-Q (Nikon) at 10× for up to 72 hours, with an interval of 7 minutes and a z-step of 7 μm before fixation.

### Follicle compression assay

A PDMS chamber was customised and fabricated with a groove with a length of 300 μm and a height of 200 μm, with a scaled 100 μm width. The PDMS chamber was mounted on the manual cell stretching system. After removing the air bubbles using the degasser, the surface of the PDMS chamber was treated with 1% Pluronic F-127 in PBS overnight to prevent the attachment and migration of TCs from the follicles onto the PDMS chamber. It was washed in PBS once, replaced with growth medium, and stretched to 50% before isolated follicles were transferred to the stretched grooves. The PDMS chamber was then relaxed gradually back to the initial width at a rate of 10 μm every 30 minutes. The follicles were cultured in the compressed state for 24 hours before fixation.

### Immunofluorescence staining

Isolated ovaries and follicles were fixed in 4% PFA at room temperature for 1 hour and 30 minutes, respectively, and washed in PBS thrice before immunostaining. Fixed tissue slices or isolated follicles were incubated in blocking and permeabilising solution (1x PBS containing 3% BSA and 0.3% Triton X-100) at room temperature for 2-4 hours, followed by incubation at 4℃ in primary antibodies diluted in the blocking solution overnight. The tissues were washed five times in washing buffer (1% BSA in 1X PBS) at room temperature and then incubated in secondary antibodies diluted in washing buffer for 4-6 hours at room temperature. They were then washed thrice in the washing buffer at room temperature before mounting. The ovarian slices were mounted into ProLong^TM^ Gold antifade mountant (P10144, Thermo Fisher) and left to cure overnight at room temperature, whereas isolated follicles were mounted into SlowFade Gold antifade mountant (S36940, Thermo Fisher) prior to imaging.

The primary antibodies used were mouse anti-YAP (H00010413-M01, Abnova, Taiwan, 1:200), rabbit anti-Ki67 (9129S, Cell Signaling Technology, 1:200), rabbit anti-collagen IV (ab19808, Abcam, 1:100), rabbit anti-laminin (nb300-144, Novus Biologicals, 1:200), rabbit anti-collagen I (ab279711, Abcam, 1:200), rabbit anti-fibronectin (ab2413, Abcam, 1:100), rabbit anti-phospho myosin light chain 2 (Ser19) (3671L, Cell Signaling Technology, USA, 1:100) and rabbit anti-NR2F2 [EPR18443] (ab211777, Abcam, 1:200). The secondary antibodies used were Alexa Fluor 488 labelled anti-mouse (A32766, Invitrogen, USA, 1:500) and Alexa Fluor 546 labelled anti-rabbit (A10040, Invitrogen, 1:500). DNA was stained with DAPI (D9542, Sigma, 2 μg/mL) and F-actin was stained with Alexa Fluor 633-labelled phalloidin (A22284, Invitrogen, 1:300). Collagen was also stained with EGFP-CNA35 (collagen-binding adhesion protein 35), which was expressed and isolated from DH5α *E. coli* with the plasmid pET28a-EGFP-CNA35. HA was stained with the Streptavidin AZDye™ 488 Monovalent Antibody Labeling Kit (ab272187, Abcam). The samples were blocked with the biotin blocking buffer for 30 minutes before incubating in the blocking and permeabilising solution. The samples were then stained with anti-hyaluronan binding protein (HABP) conjugated to biotin (AMS.HKD-BC41, Amsbio, 1:200) and Streptavidin AZDye™ 488 (Monovalent) labeling reagent, respectively.

For fixed cells on PA and PDMS hydrogels, they were incubated using the blocking and permeabilising solution for 10 minutes and blocked using washing buffer for one hour at room temperature. The TCs were then stained with primary antibodies in 1% BSA in PBS overnight at 4℃. Afterwards, the TCs were washed thrice with PBS and stained with secondary antibodies in washing buffer overnight at 4℃. After washing thrice with PBS, the PA and PDMS hydrogels were mounted onto glass slides with antifade mountant, overnight at room temperature.

### EdU incorporation assay

Follicles were incubated with 10 µM EdU (EdU Staining Proliferation Kit, abcam, ab219801) for 30 mins under optimal growth conditions. They were washed with WB and were fixed and permeabilised. The EdU reaction solution was prepared as per the manufacturer’s specifications. The follicles were incubated in the reaction solution for 2 hours at RT in the dark. They were washed and incubated with DAPI for 2 hours at RT in the dark before being washed and then mounted for imaging.

### Confocal imaging

All fixed samples were imaged with the Nikon A1Rsi confocal microscope. The acquisition was managed by NIS Elements in a 16-bit 1024×1024 format, using lasers 405 nm, 488 nm, 546 nm, and 640 nm. Samples were imaged using Apo WI λS DIC N2 objectives at various magnifications depending on the protein of interest, and samples were stitched with a 10% overlap at high magnification (100×).

### Testosterone enzyme-linked immunosorbent assay

Isolated TCs on PA gels with different substrate stiffness were cultured with 1.5 IU/mL hCG for 3 days. The culture medium was aspirated for testosterone concentration measurement using the mouse Testosterone enzyme-linked immunosorbent assay (ELISA) kit (FineTest, EM1850), whereas the cultured TCs were fixed and used for cell density quantification. The culture medium was serially diluted to ensure that the protein concentration was within the detectable range of the ELISA kit. The ELISA plate was washed twice with the washing Buffer before adding the standards and samples into the wells. The biotin-labelled antibody solution was added and mixed well by gently shaking and incubated at 37℃ for 45 minutes. The plate was washed thrice, and HRP-Streptavidin Conjugate (SABC) was added and incubated at 37℃ for 30 minutes. The plate was washed five times before adding the TMB substrate solution and incubated at 37℃ for 10-20 minutes. The stop solution was added, and the samples were measured using a plate reader at 450 nm.

### Data quantification and analysis

#### Quantification of TC YAP N/C ratio

The nuclei and cytoplasm were identified using DAPI and phalloidin, respectively. For cells, the maximum projection of the images was obtained using FIJI and its Z Project function. For tissue sections and isolated follicles, the equatorial plane of the follicles was used for analysis. For compressed follicles, multiple planes of the follicles were used for analysis. The cell nuclei and area of the TCs were then drawn manually and used to measure the YAP intensities in the nucleus and cytoplasm, respectively. The YAP nuclear to cytoplasmic ratio was then calculated for each cell; values were averaged for each follicle and plotted. For *in vitro* experiments, YAP N/C ratio was averaged for every imaging field of view and plotted.

### Quantification of TC proliferation index

The total number of cells was determined using DAPI, whereas the number of proliferative cells was measured using the number of Ki67-positive cells, with Ki67 as a binary marker. The proliferation rate was calculated by dividing the Ki67-positive count by the DAPI count.

### Quantification of BM intensity and thickness

A line with a width of 3 pixels was drawn on top of the BM and the intensities of collagen IV and laminin were measured using Fiji. The values were divided by the intensity at the oocyte and termed as the normalised matrix intensity. For quantifying BM thickness, 20 linescans were drawn perpendicular to the BM for each follicle in the SEM images at the equatorial plane using FIJI. The width and standard deviation of each linescan was measured; the average calculated and termed as the BM thickness in each follicle.

### Quantification of BM elasticity

AFM measurement curves were analysed using the JPKSPM Data Processing software by the batch processing function. Since the first 70 nm (indicative of the measured BM thickness) of the curve consists of relatively few data points, fitting within this regime is prone to noise and results in large fitting errors. We therefore fitted the first 150 nm of the curve to the Hertz/Sneddon model, and the average values were used to calculate the Young’s modulus of the BM elasticity and standard deviation. The follicles used for AFM, both control and collagenase IV-treated, were fixed and stained with DAPI and CNA35 to validate the absence of TCs and accurate measurement of the BM elasticity.

### Quantification of ECM and pMLC intensity at TCs

The boundary of the basal TC layers was determined using phalloidin. For bulk basal TCs comparison, the basal TC layer and oocyte area were drawn manually using FIJI. The intensities of collagen I, HABP and pMLC were measured using the basal TC layer ROI, divided by the intensity at the oocyte and termed as normalised intensity. To measure high intensity pixels from the stacks, a max projection of the HABP channel was calculated in every follicle. All pixels with intensity beyond 75% of the maximum intensity across the entire volume was measured. Follicles of similar size and with the same z-planes and acquisition settings were used for this analysis.

### Quantification of HA-other ECM co-localisation

For the co-localisation study, the HABP was co-stained with either collagen IV or fibronectin. A line scan was drawn across the BM, from the intrafollicular region outwards to the basal TCs using Fiji. The intensity profile was plotted using OriginPro, and the intensity peaks were extracted. The difference in intensity peaks between HABP and collagen IV or fibronectin was calculated and plotted.

### Quantification of PA gel stiffness

The Young’s modulus of the PA gels was calculated by fitting the loading portion of the force-displacement curve to the Hertzian contact model using the ‘Prova’ (Optics11Life, Amsterdam, The Netherlands) software, assuming a Poisson’s ratio ν of 0.5. A total of four independent hydrogel samples were analysed per experimental group.

### Quantification of TC mitotic occurrence and timescale *ex vivo*

The holotomography images imaged at various planes of the cultured follicles was used to count for any mitotic event occurring at the theca cells over 4 hours. The average mitotic occurrence in each follicle was calculated and plotted in control and treated samples. The time between each theca cell from the start of rounding to separating into two daughter cells is measured and termed as TC division time.

### Quantification of TC cell division and motility *in vitro*

Isolated TCs were cultured on TMSPMA-treated coverslips to confluency and imaged live via Biostation IM-Q. TC division events were counted and normalised by the cell density and sample area, for 2 hours before and after pharmacological perturbation. For TC motility, particle image velocimetry (PIV) analysis was performed using PIVlab, a MATLAB graphical user interface (GUI), 4 hours before and after pharmacological perturbation. For the PIVlab analysis, three band passes with interrogation areas and steps of 128 x 64, 64 x 32 and 32 x 16 pixels were used. The thresholds used for the filter low contrast and filter bright object were 0.01 and 0.85, respectively. At the end of the pipeline, the median velocity of the TCs was then extracted via MATLAB and plotted via OriginPro.

### Purity assessment of isolated TCs

The number of NR2F2-positive nuclei was divided by the total number of DAPI stained nuclei in each image for both GCs and TCs to calculate NR2F2-positive cells. ALP-positive cells were counted by dividing the number of ALP-stained cells with the total number of cells in each image. The cell boundaries of both cell types were drawn manually using FIJI and the circularity was measured using the cell boundaries with the Shape descriptors option in Fiji.

### Quantification of TC tissue fraction on hemicylindrical substrates

Isolated TCs were cultured on hemicylindrical substrates and imaged live via Biostation IM-Q. The area occupied by the cells on the hills or valleys was manually annotated and measured using Fiji, at two time points, 0 and 60 hours. The tissue fraction of the TCs was calculated by dividing the TC-covered area by the total area of the hills or valleys, before and after 60 hours.

### Quantification of testosterone secretion by ELISA

A standard curve is plotted using the intensity values measured from the standards using OriginPro 2021b. The testosterone concentration for each sample is determined using the standard curve and normalised by the cell density. The normalised fold change in testosterone concentration is calculated by dividing the testosterone concentration by the average testosterone concentration of TCs in Fig. S4F; and by the average testosterone concentration at 19.2 kPa in Fig. S6C.

### Transcriptomic analysis of published dataset

scRNA-seq data from the ovaries of postnatal day 21 mice generated using the 10× Chromium platform was obtained from GSA:CRA010027 (CRR703610), GSA:CRA012042 (CRR838123) and GEO:GSE268466 (GSM8291634). Raw FASTQ files were aligned using Cellranger (v10.0.0, 10x Genomics) with GRCm39 reference genome. Downstream analyses were performed using Seurat (v5.3.1). Cells with fewer than 200 detected genes or with mitochondrial RNA content exceeding 20% were excluded from analysis. scRNA-seq datasets were then normalised with SCTransform and integrated using Harmony (86). Cell clusters were isolated and subclustered according to Morris et al (87).

### Quantification of follicle growth

Follicle diameters *D* were measured by taking the average of the major and minor axes from the Fit Ellipse tool in ImageJ. Growth rate *G*(*r*) of follicles was defined as:

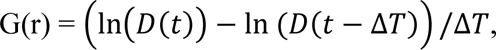

where Δ*T* is the time interval between two consecutive time points and Δ*T* is 1 day. The growth rate for diameters >180 µm was used to plot G(r) at the maturation phase.

### Statistics

All graphs and statistical tests were created using OriginPro. No statistical method was used to predetermine the sample size. *n* represents the number of tissues/follicles/cells in the representative data shown in the figures. Sample sizes were determined based on previous studies in similar fields. The data were tested for significance using two-tailed Mann-Whitney U test or one-sample t-test. Boxplots show the mean (star), median (centre line), quartiles (box limits) and 1.5x interquartile range (whiskers). No specific methods were used for random allocation of samples into groups. All experiments were performed on at least four biological replicates.

