## Supplementary Information for "Theca cell mechanosensing and regulation of follicular extracellular matrix during ovarian follicle development"

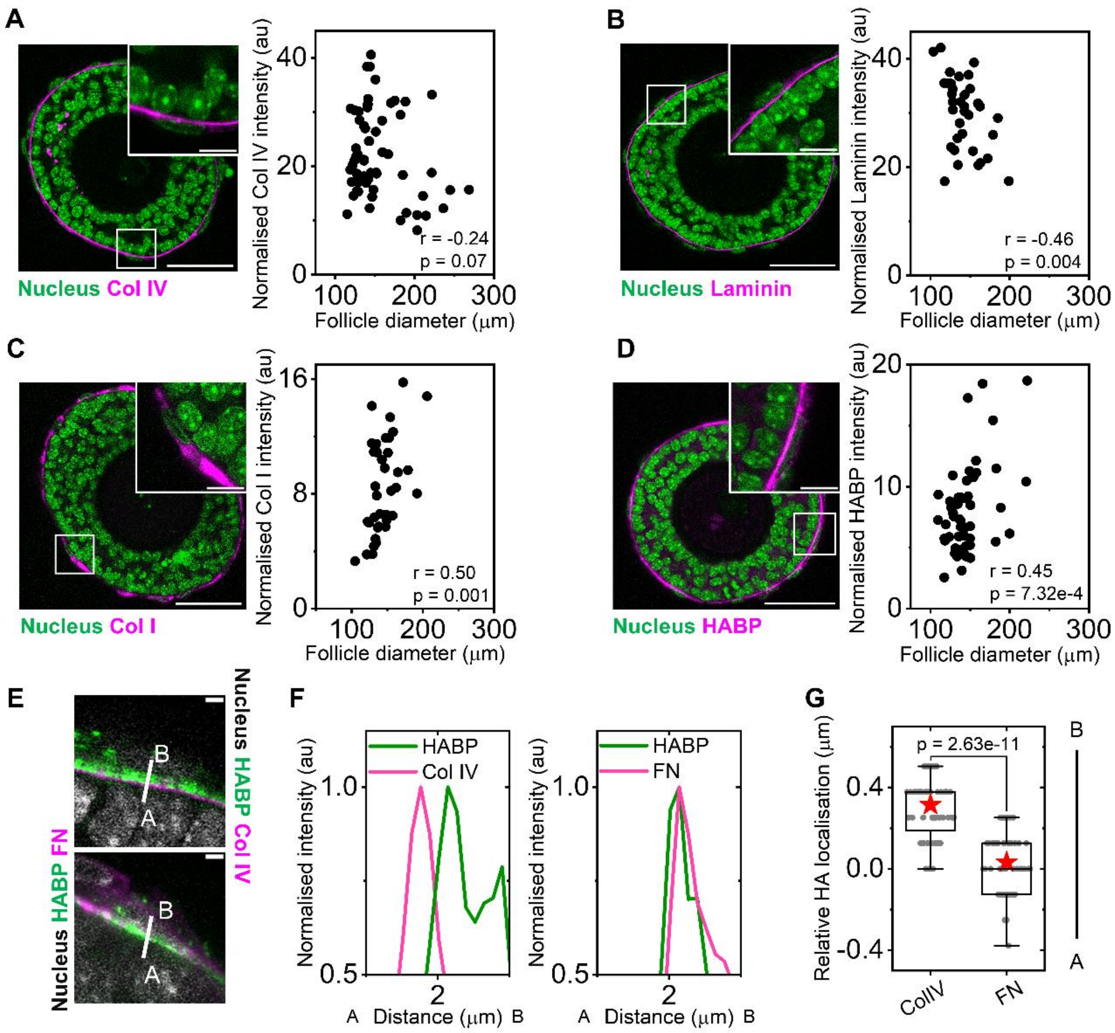


**Fig. S1.** ***Ex vivo*** **characterisation of TC microenvironment during secondary follicle development.** (A) Left: representative image of an isolated secondary follicle stained for DAPI (green) and collagen IV (magenta). Scale bar: 50 µm. Inset: zoomed-in section of the overlaid white box. Scale bar: 10 µm. Right: scatter plot of collagen IV intensity against follicle diameter. $n$ = 59 follicles. (B) Left: representative image of an isolated secondary follicle stained for DAPI (green) and laminin (magenta). Scale bar: 50 µm. Inset: zoomed-in section of the overlaid white box. Scale bar: 10 µm. Right: scatter plot of laminin intensity against follicle diameter. $n$ = 36 follicles. (C) Left: representative image of an isolated follicle stained for DAPI (green) and collagen I (magenta). Scale bar: 50 µm. Inset: zoomed-in section of the overlaid white box. Scale bar: 10 µm. Right: scatter plot of normalised collagen I intensity at the TCs against follicle diameter. $n$ = 38 follicles. (D) Left: representative image of an isolated secondary follicle stained for DAPI (green) and HABP (magenta). Scale bar: 50 µm. Inset: zoomed-in section of the overlaid white box. Scale bar: 10 µm. Right: scatter plot of normalised HABP intensity at the TCs against follicle diameter. $n$ = 52 follicles. (E) Top: zoomed-in image of an isolated follicle stained with DAPI (grey), HABP (green) and collagen IV (magenta). Bottom: zoomed-in image of isolated follicle stained with DAPI (grey), HABP (green) and fibronectin (magenta). Scale bar: 2 µm. (F) Left: intensity profile of HABP and collagen IV for the line scan marked in white (top, Supp. Fig. 1E). Right: intensity profile of HABP and fibronectin for the line scan marked in white (bottom, Supp. Fig. 1E). (G) Boxplots of HA localisation relative to collagen IV and fibronectin respectively. $n$ = 16 follicles each; 3 line-scans in each follicle. Significance was determined by two-tailed Mann-Whitney U test (pairwise) in G. Boxplots show the mean (star), median (centre line), quartiles (box limits) and 1.5x interquartile range (whiskers). Pearson correlation coefficient ($r$) and significance ($p$, two-tailed test) are noted in the plots for A-D. All data are from at least three biological replicates.

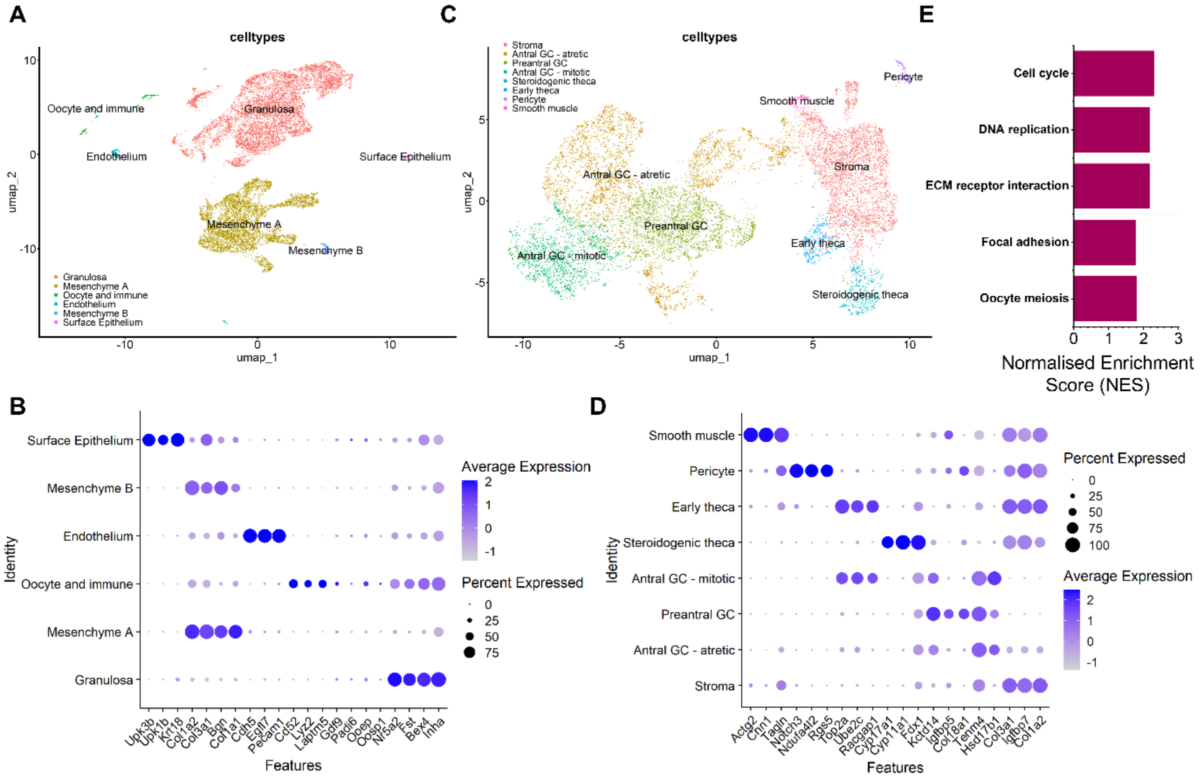


**Fig. S2. Transcriptomic analysis of early theca cells in follicles from three-weeks old mice.** (A) UMAP plot showing clustering of all major cell types identified in the ovary. (B) Corresponding dot plot displaying the expression of key marker genes defining each cell type. Dot size represents the percentage of cells expressing the gene, and colour intensity indicates the average expression level. (C) UMAP plot showing clustering of all major cell types identified in the mesenchyme clusters. (D) Corresponding dot plot displaying the expression of key marker genes defining each cell type. Dot size represents the percentage of cells expressing the gene, and colour intensity indicates the average expression level. (E) Bar plots of normalized enrichment scores (NES) for significantly enriched KEGG pathways for *Mus musculus* identified via GSEA in early theca cells vs. pre-antral granulosa cells. Magenta bars represent positive NES with adjusted p<0.05. All p-values by Wilcoxon rank-sum test.

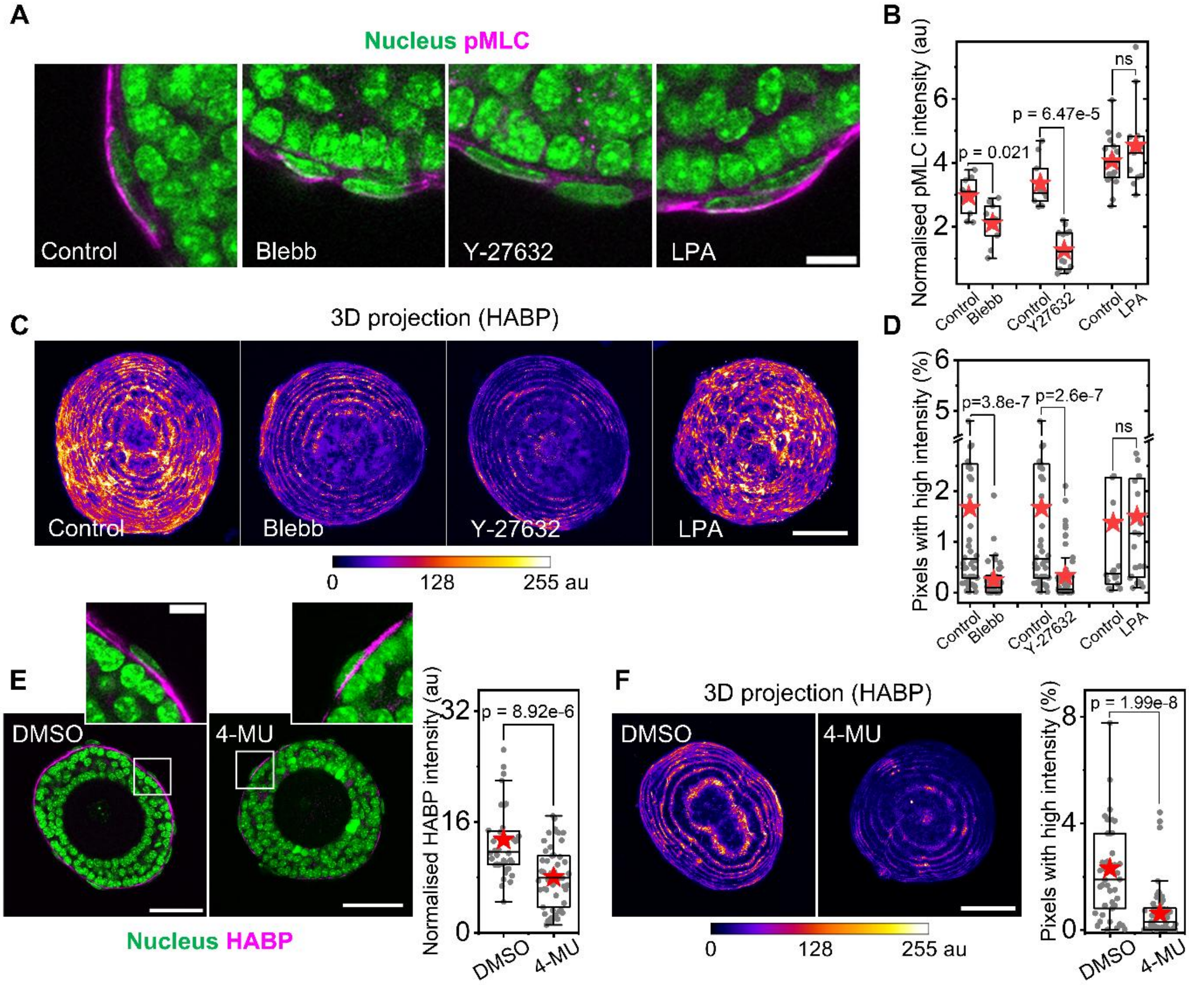


**Fig. S3. Validation of pharmacological perturbations targeting HA synthesis and TC contractility *ex vivo*.** (A) Representative images of zoomed-in sections of isolated follicles stained for DAPI (green) and pMLC (magenta) under different contractility perturbations. Scale bar: 10 µm. (B) Boxplots of normalised pMLC intensity under different contractility perturbations. $n$ = 46 (control), 40 (blebb); 46 (control), 49 (Y-27632); 18 (control), 11 (LPA) follicles. (C) Representative images of maximum intensity 3D projections for HABP of follicles in the different conditions. Scale bar: 50 µm. (D) Boxplots of pixels with high HABP intensity in follicles under contractility perturbations. $n$ = 46 (control), 40 (blebb); 46 (control), 49 (Y-27632); 18 (control), 24 (LPA) follicles. (E) Left: representative images of isolated follicles stained for DAPI (green) and HABP (magenta) on inhibiting HA synthesis. Scale bar: 50 µm. Inset shows zoomed-in sections of the overlaid white boxes. Scale bar: 10 µm. Right: boxplots of normalised HABP intensity of basal TCs during inhibition of HA synthesis. $n$ = 40 (DMSO), 51 (4-MU) follicles. (F) Left: representative images of maximum intensity 3D projections for HABP of follicles in the different conditions. Scale bar: 50 µm. Right: boxplots of pixels with high HABP intensity in follicles under contractility perturbations. $n$ = 42 (DMSO), 67 (4-MU) follicles. Significance was determined by two-tailed Mann-Whitney U test (pairwise) in C, E. Boxplots show the mean (star), median (centre line), quartiles (box limits) and 1.5x interquartile range (whiskers). All data are from at least three biological replicates.

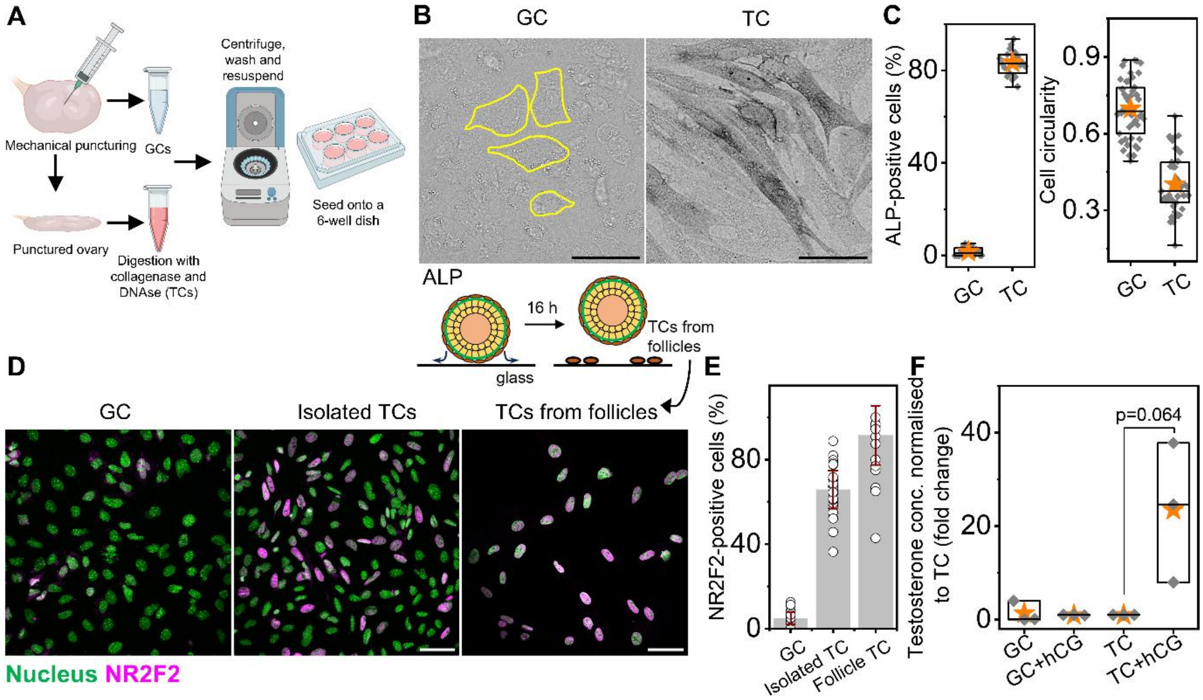


**Fig. S4. Purity assessment of primary TC isolation.** (A) Schematic of protocol to isolate TCs and GCs from bulk ovaries. The schematic is made using BioRender. (B) Representative images of isolated GCs and TCs stained for alkaline phosphatase (ALP). The GCs were outlined (yellow) due to the lack of ALP staining. Scale bar: 50 µm. (C) Left: boxplots of cell circularity of isolated GCs and TCs. $n$ = 45 (GCs), 43 (TCs) cells. Right: boxplots of ALP-positive cells (%) from the isolated GCs and TCs. $n$ = 18 images. (D) Representative images of isolated GCs and TCs using the method mentioned above; and of cells migrating from the follicles onto substrates stained with DAPI (green) and NR2F2 (magenta). Scale bar: 50 µm. Schematic of TCs retrieved from follicles is shown on top. (E) Boxplots of NR2F2-positive cells (%) in the different conditions. Bars and error bars represent the mean and standard deviation. $n$ = 30 images. (F) Plot of testosterone concentration secreted by culturing isolated GCs and TCs, without or with hCG. N = 3. Significance was determined by two-tailed Mann-Whitney U test (pairwise) in C, F. Boxplots show the mean (star), median (centre line), quartiles (box limits) and 1.5x interquartile range (whiskers). All data are from at least three biological replicates.

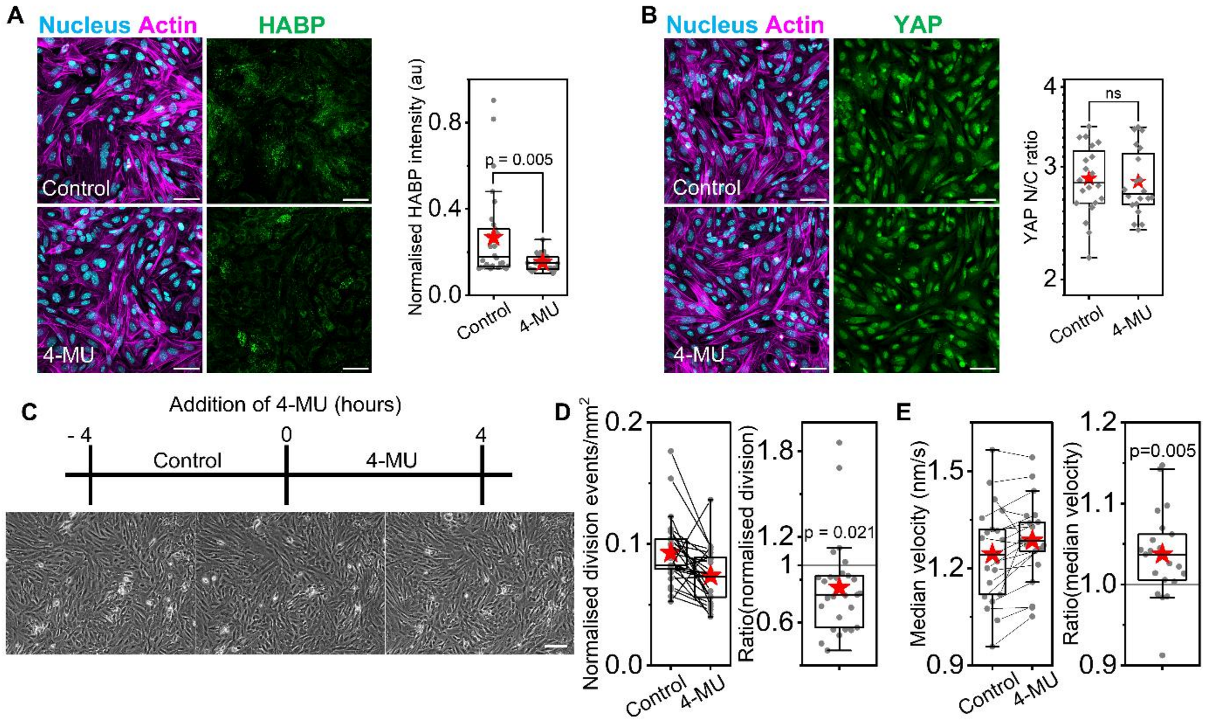


**Fig. S5. Inhibition of HA synthesis impacts TC functions *in vitro*.** (A) Left: representative images of isolated TCs cultured on glass stained for DAPI (cyan), phalloidin (magenta) and HABP (green) upon inhibition of HA synthesis. Scale bar: 50 µm. Right: boxplots of normalised HABP intensity of TCs on 4-MU treatment. $n = 30$ images. (B) Left: representative images of isolated TCs cultured on glass stained for DAPI (cyan), phalloidin (magenta), and YAP (green) upon inhibition of HA synthesis. Scale bar: 50 µm. Right: boxplots of YAP N/C ratio (in log scale) of TCs on 4-MU treatment. $n = 20$ images. (C) Representative images of TC dynamics before and after 4-MU treatment. Scale bar: 100 µm. (D) Left: boxplots of normalised cell division events upon inhibition of HA synthesis. Right: boxplot of normalised cell division events after 4-MU treatment. $n = 27$. (E) Left: boxplots of median velocity upon inhibition of HA synthesis. Right: boxplot of relative change in TC median velocity after 4-MU treatment. $n = 21$. Significance was determined by two-tailed Mann-Whitney U test (pairwise) in A, B; and one-sample t-test (test mean = 1) in D, E. Boxplots show the mean (star), median (centre line), quartiles (box limits) and 1.5x interquartile range (whiskers). All data are from at least three biological replicates.

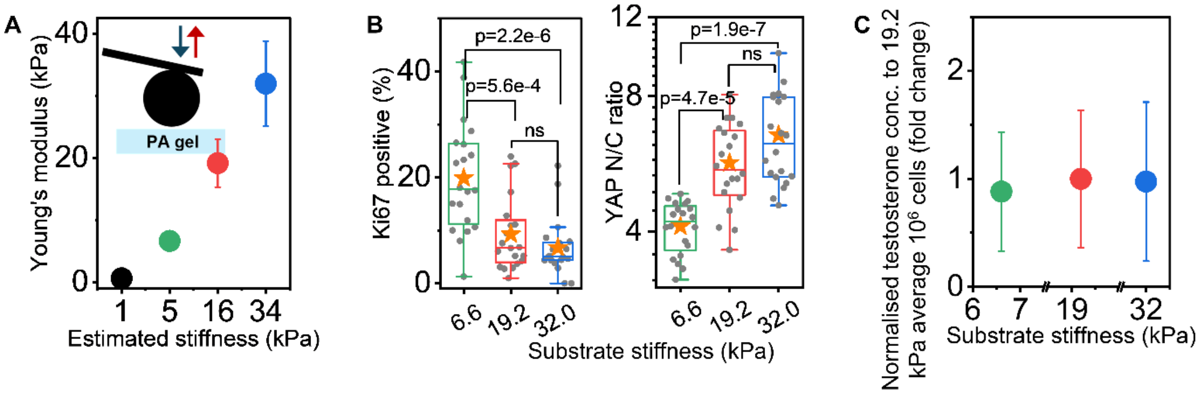


**Fig. S6. TC YAP signalling remains consistent on confluency and upon hCG induction.** (A) Plot of the average Young's modulus measured for different polyacrylamide gel stiffness conditions. $N = 4$. Inset: schematic represents how nanoindentation is used for measurements on prepared PA gels. (B) Left: boxplots of Ki67-positive TCs cultured in high cell density on the varying substrate stiffness, with the addition of hCG. $n = 20$. Right: Boxplots of YAP N/C ratio (in log scale) of TCs cultured in high cell density on varying substrate stiffness, with the addition of hCG. $n = 300$ cells. (C) Plot of normalised testosterone concentration secreted by TCs cultured on varying substrate stiffness, with the addition of hCG. $n = 8$. Significance was determined by two-tailed Mann-Whitney U test (pairwise) in B. Boxplots show the mean (star), median (centre line), quartiles (box limits) and 1.5x interquartile range (whiskers). All data are from at least three biological replicates.

### Supplementary Movies

**Movie S1:** Timelapse imaging of a secondary follicle using holotomography. White box overlaid on the movie shows a TC undergoing division and is represented in Fig. 1J (top). Scale bar: 50 µm.

**Movie S2:** Timelapse imaging of a follicle from H2B-mCherry transgenic mouse using fluorescence imaging. White box overlaid on the movie shows a TC undergoing division and is represented in Fig. 1J (bottom). Scale bar: 50 µm.

**Movie S3:** TC division dynamics captured by holotomography imaging at zoomed-in sections of a follicle in control (left) and 4-MU (right) treatment. Scale bar: 10 µm.

**Movie S4:** Timelapse imaging captured by brightfield imaging of isolated TCs on glass before (control, left) and after (right) adding 4-MU. Scale bar: 100 µm.

**Movie S5:** TC dynamics on regions of positive curvature (hills) on hemicylindrical substrates captured by brightfield imaging. Scale bar: 100 µm.

**Movie S6:** TC dynamics on regions of negative curvature (valleys) on hemicylindrical substrates captured by brightfield imaging. Scale bar: 100 µm.
